# Limits to resilience of Afroalpine vegetation to grazing and burning: a case study of grasses from the Drakensberg Mountain Centre, southern Africa

**DOI:** 10.1101/2021.04.06.438591

**Authors:** Steven P. Sylvester, Robert J. Soreng, Aluoneswi C. Mashau, Mitsy D.P.V. Sylvester, Anthony Mapaura, Vincent Ralph Clark

## Abstract

1. High-elevation Afroalpine ecosystems of the Drakensberg Mountain Centre (DMC) of Lesotho and South Africa, renowned for their high endemism and key ecosystem services, are socio-ecological systems that have seen human activity for millennia. However, their responses to land management practices are understudied. Controversy over their natural state has also led to conflicting policies and management emphases.
2. Focusing on the crucial ecosystem-modulating component, grasses (Poaceae), we evaluate the response of DMC Afroalpine vegetation to human impact through grazing and burning. Grass species associations were recorded from grassland, shrubland and wetland-riparian-seep ecotypes across a range of grazing and fire regimes to document relationships between abiotic conditions, disturbance, and taxonomic diversity and composition.
3. CCA of grass community composition retrieved a large cluster of plots of mixed grazing and burning regimes with no particular environmental vector correlated with them. Other smaller groups of plots separated from these were associated to heavy grazing, bioclimatic variables, slope gradient, and aspect. Indicator species analyses found DMC endemic grasses were associated to low grazing, while alien grasses were associated to heavy grazing. GLMs found little difference between ecotype-disturbance categories with regards plant species richness, mean alpha hull=2 range-size of native and sub-Saharan endemic grasses, and site-level Sørensen beta diversity (βsor). Some differences were noted, including the highest cover and proportion of DMC endemics being found in low-grazed grassland, and highest cover and proportion of alien grasses and highest plot-level βsor being found in heavily grazed ecotypes. Relative importance analyses found grazing regime to be the main influence on cover and proportion of DMC endemic and alien grasses. Partial Mantel tests found mean annual temperature and grazing regime to be the main influence on plot-level βsor.
4. Synthesis: Taxonomic diversity and composition of DMC Afroalpine grasslands was relatively unaffected by moderate grazing and intense burning, although heavy grazing had a largely detrimental impact, with its ubiquity across the DMC a major cause for concern. High levels of endemism, coupled with the above data emphasizing the robustness of DMC grasslands to disturbance, also supports Afroalpine grasslands as a natural component of the DMC. This research reinforces the natural grass-dominated nature of the DMC as a social-ecological system where sustainable management is possible thanks to its resilience to grazing and burning, although current widespread overgrazing requires urgent attention.

## 1 Introduction

The grass-dominated Drakensberg Mountain Centre (DMC; Carbutt, 2019) is the highest elevation mountain system in southern Africa and represents the core alti-montane (Afroalpine and Afromontane) grassland in southern Africa (White, 1983). These alti-montane grasslands provide important ecosystem services (Taylor et al., 2016; Ngwenya et al., 2019) and are considered a continental hotspot of botanical diversity and endemism (Carbutt & Edwards, 2004; Brand et al., 2019; Carbutt, 2019) with c. 9% of angiosperm species endemic to the DMC (Carbutt, 2019). This high endemism may relate to their isolation, being found 2800 km south from the closest Afroalpine ecosystems in Central and Eastern Africa (Carbutt, 2020) as well as an obviously differing climate regimen from tropical Afroalpine systems (Carbutt & Edwards, 2015). The vegetation structure of the DMC differs markedly from Afroalpine areas of tropical Africa in the absence of a physical treeline and the dominance of an exceptionally rich grassland flora and fauna (as opposed to forest) with very strong biogeographical connections to the Fynbos Biome of the Cape (Meadows & Linder, 1993; Mucina & Rutherford, 2006; Gehrke & Linder, 2014; Carbutt & Edwards, 2015; Carbutt, 2019). DMC ecosystems are described as mosaics of forest ‘islands’ among a ‘sea’ of grassland or heathland, with controversy surrounding whether these patterns are a result of long-term human impact or whether these grasslands are indeed natural (Meadows & Linder, 1993; Adie et al., 2017). Indeed, humans are reported to have been present at high elevations in the DMC as far back as c. 80,000 years ago (Stewart et al. 2016). Because these high-elevation grasslands have been exposed to human activity for many millennia, they can be considered socio-ecological systems (SES), but anthropogenic pressures have changed significantly over the last decades, mainly with regards to grazing and burning (Carbutt, 2020).

The DMC grasslands support a large agrarian community, with subsistence-based livestock herding the dominant land-use occupying 79% of the land (Adelabu et al., 2020). However, current land management practices are considered unsustainable and could lead to severe habitat degradation, which also compromises other ecosystem services (carbon sequestration, water production, etc.) and entrenches poverty cycles (Adelabu et al., 2020; Carbutt, 2020; Turpie et al., 2021). Added to this is the threat of global warming, with projected range contraction of montane vegetation to higher elevations and subsequent biodiversity loss (Bentley et al., 2019). Despite these threats, Afroalpine grasslands of the DMC remain understudied regarding their response to human influence, with most ecological, paleoecological and floristic research focused on lower-elevation Afromontane grasslands of southern Africa (e.g. Joubert et al., 2017; Lodder et al., 2018; Breman et al., 2019; Morris et al., 2021) and Afroalpine grasslands found in East and Central Africa (e.g. Gehrke & Linder, 2014; Johansson et al., 2018; Lézine et al., 2019; Vidal & Clark, 2020). A recent overview of all Afroalpine vegetation (Carbutt, 2020) found it to be currently under significant threat from human-induced land transformation, degradation and unsustainable use and recommended large-scale policy shifts. Controversy exists, however, over how human impact alters the dynamics of these ecosystems (e.g. Meadows & Linder, 1993) and whether human management of fire and grazing positively influences diversity and ecosystem services, with little empirical data available at landscape scale on the complex relationships between fire, grazing, biodiversity, and ecosystem function. This information is necessary to determine the degree to which these unique ecosystems are at risk while feeding back to ongoing debates regarding how ‘natural’ these ecosystems are and the true extent of human impact in these landscapes. This is particularly important for influencing conservation and management policies, with changing governmental policies (e.g. increasing carbon storage via fire suppression, reforestation) not based on scientifically sound information potentially causing tremendous ecological damage (Johansson et al., 2018; Bond et al., 2019).

In this paper, we focus on grasses (Poaceae) as a study system to improve functional understanding of Afroalpine grasslands of southern Africa. Grasses are unique in the plant kingdom: they build and modulate open canopy ecosystems covering more than 25% of dry land across the globe. Grass-dominated ecosystems differ between climatically equivalent regions, partly due to traits of the locally dominant grass species (e.g. Lehmann et al., 2011; Strömberg, 2011). Grasses are also challenging to identify and classify, and are here studied by dedicated specialists. Using grasses as a study system, the objectives of this paper are to assess the influence of grazing and burning regimes, plus other abiotic variables, on taxonomic diversity and composition (including endemicity, range size rarity, and alien components) of DMC Afroalpine vegetation.

## 2 Materials and Methods

### 2.1 Study area

The study was conducted within Afroalpine grasslands of the DMC alpine sub-centre of Lesotho and South Africa (Fig. 1) (the former Drakensberg Alpine Centre of Van Wyk & Smith, 2001, and Carbutt & Edwards, 2004, 2006) which are dominated by C_3_ grass species (Killick, 1994; Bentley & O’Connor, 2018; Brand et al., 2019). The study was conducted between 1 Feb. and 26 Mar., 2020. Ten main field study sites were selected from throughout the DMC and included: Sani Pass area (KwaZulu-Natal, South Africa; adjacent Lesotho); Witsieshoek and Mont-aux-Sources (Free State & KwaZulu-Natal, South Africa; adjacent Lesotho); Tiffindell Ski resort and surroundings, Naudes Nek, and Bastervoetpad Pass areas (Eastern Cape, South Africa); Matebeng Pass, Sehlabathebe National Park, AfriSki Ski Resort and surroundings, and Bokong Nature Reserve (Lesotho). These sites represent the main ecological variations across the DMC Afroalpine zone. Although certain literature states the Afroalpine zone to be >2800 m alt. (e.g. Carbutt, 2020), we chose to define this based on the dominant cover (>50% cover) of grasses being C_3_ grass species, with certain lower-elevation areas in the Eastern Cape having cooler and wetter microclimates similar to those of higher elevations and which were dominated by C_3_ grass species. Elevations ranged from 2235–3292 m alt., and were generally >2600 m alt. (mean= 2894.9 m alt.; standard deviation= 234.1). These areas were geologically dominated by continental flood basalt lavas of the Drakensberg Group (Karoo Supergroup), dated as Early Jurassic (ca. 183 Ma) (Knight & Grab, 2015).

**Figure 1.**
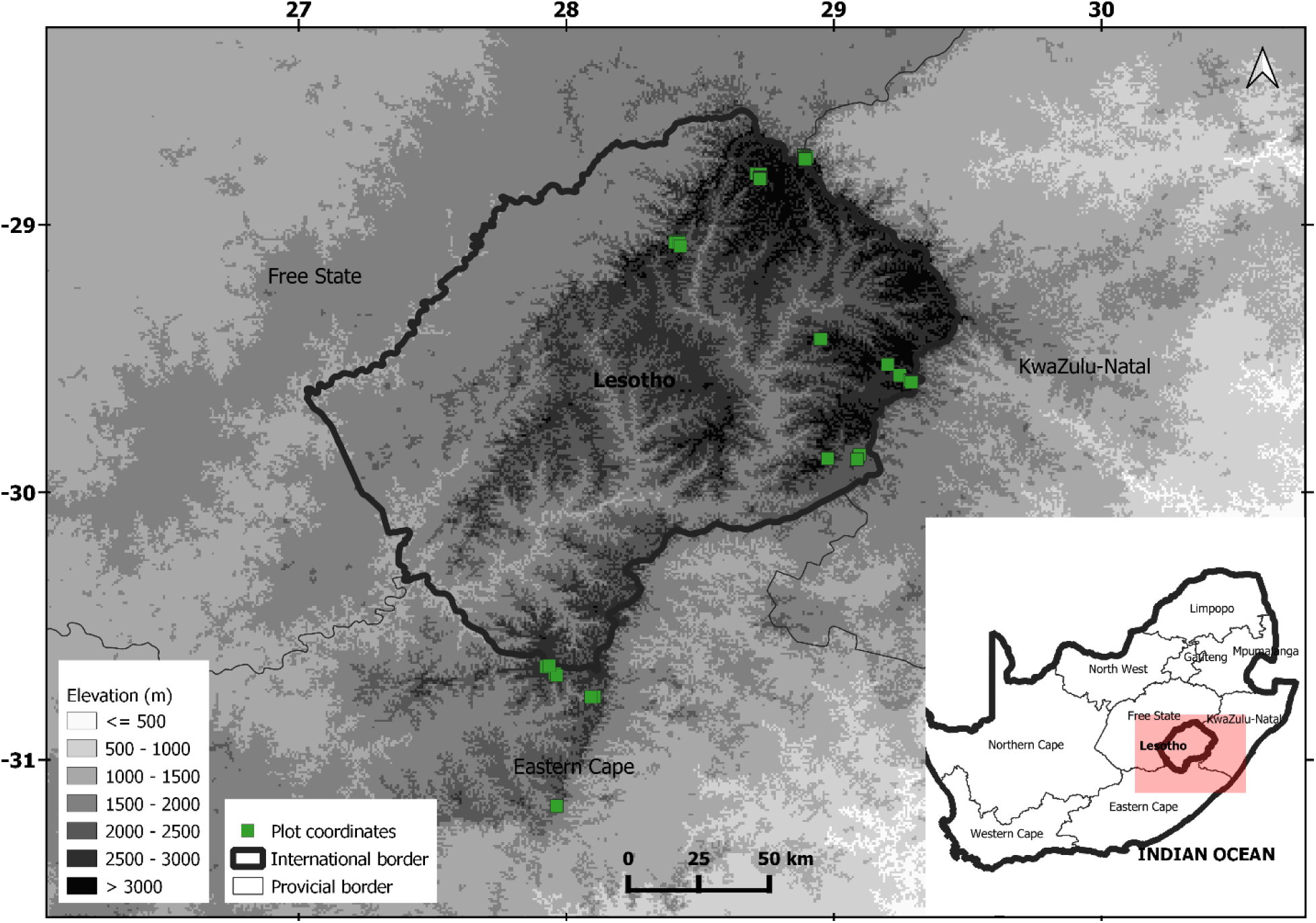
Map of the study sites across the DMC.

### 2.2 Sampling design

A total of 222 2×2m quadrat plots were studied from 10 main sites across the DMC (Fig. 1). Sites were divided into three ecotypes: grassland, shrubland (where woody phanerophytes dominated the area and covered >15% within quadrats), and wet areas (including seeps, streamside, riverside, or wetland). Ecotypes were further divided into areas that experience differing grazing and burning regimes, with plots assigned to three broad disturbance categories, i.e. low, moderate, or high, based on informal discussions with local people and within-site observations. None of the sites could be described as containing pristine vegetation, although areas of low grazing and low burning were usually found within protected areas which could hypothetically represent a more natural state. Grazing regimes were categorized according to a combination of yearly and long-term stocking densities (obtained via informal discussions with locals), dung presence, observed disturbance by grazing animals, distance to settlements or ranches, and distance to paths. A ‘low’ grazing regime was characterized by low stocking densities (<1 animal/ha), little or no observed dung and animal disturbance in the area (usually <5% quadrat cover), a proximity to paths of generally >30 m, and a proximity to settlements of generally >3 km. A ‘moderate’ grazing regime was characterized by medium stocking densities (1–2 animals/ha), dung and animal disturbance observed in the area (usually 25–60% quadrat cover noted to be grazed or trampled), a proximity to paths of generally <30 m, and a proximity to settlements of generally <1 km. A ‘high’ grazing regime was characterized by high stocking densities (>2 animals/ha), very notable dung and animal disturbance observed in the area (usually >80% quadrat cover noted to be grazed or trampled), a proximity to paths of generally <5 m, and a proximity to settlements of generally <0.1 km. Burning regimes were categorized according to the return interval (RI) between fires, i.e. low = >3 years RI, moderate = 2–3 years RI, high = ≤1 RI. Not all sites contained the full combination of ecotypes and disturbance regimes, and across all sites there was a total of 22 ecotype-disturbance categories. At least three quadrats were studied per ecotype-disturbance category, with the number of quadrats depending on the size and heterogeneity of the sites in terms of elevational range, slope gradient and aspect, zonality, as well as time permitted to study individual areas. Quadrats were spaced with at least 5 m distance between them to sample as much variability in the vegetation as possible. For each quadrat, the following were recorded: grass species cover and mean vegetative height; overall vascular plant species richness; woody phanerophyte cover and mean height; % bare ground; % rock; % litter; slope inclination and aspect; % disturbance by grazing and burning; with other metadata also sampled e.g. elevation and coordinates taken using GPS. This approach is methodologically comparable to previous studies of high-elevation grassland ecosystems of the Andes (Sylvester et al., 2014, 2017).

A representative herbarium sample of each grass species was made for each site or region, depending on the morphological variability and phylogenetic complexity of the taxa, and DNA vouchers were also taken for phylogenetically complex taxa or morpho-species of interest. *In situ* photographs were also taken. Voucher specimens were given to the National Herbarium of South Africa (PRE), the Bews Herbarium (NU), and the United States National Herbarium (US; received Jan. 4 2021).

### 2.3 Environmental variables

Aside from plot metadata recorded *in situ* as mentioned above, 19 bioclimatic (BioClim) variables (Table S1), with a resolution of 30 seconds (∼1 km^2^), were also extracted from WorldClim (www.worldclim.org) and recorded for each quadrat.

### 2.4 Range size calculation

We extracted and analysed all occurrence records of indigenous southern African Poaceae from herbarium specimens housed in the National Herbarium (PRE), Pretoria; Compton Herbarium (NBG and SAM), Cape Town; KwaZulu-Natal Herbarium (NH), Durban); herbarium acronyms following Index Herbariorum (Thiers, 2021). All the above-mentioned herbaria are managed by the South African National Biodiversity Institute (SANBI), with information held in the BODATSA/BRAHMS database (http://newposa.sanbi.org/, accessed 13/12/2019). Geo-referenced data from Fish et al. (2015) and the BODATSA/BRAHMS database were augmented with location data from specimens collected by Steven P. Sylvester et al.. Indigenous species found outside of the FSA region (Fish et al., 2015) had specimen records augmented with those collated from GBIF (https://www.gbif.org/; accessed 11/12/2020). Specimen records of *Lachnagrostis lachnantha* (Nees) Rúgolo & A.M. Molina and *Themeda triandra* Forssk. were also augmented with georeferenced specimens from the escarpment and the high plateau areas of Yemen based on Wood (1997). *Koeleria capensis* (Steud.) Nees, although stated by Plants of the World Online (2021) to occur in Yemen based on Wood (1997), was restricted in these analyses to sub-Saharan Africa due to Yemen specimens being redetermined as *K. macrantha* (Ledeb.) Schult. by Tom Cope in 2007 (John R.I. Wood, pers. communication).

The occurrence data were cleaned by removing all records with the following issues: no geographical coordinates, duplicates, localities in the sea or other waterbodies, country centroids and localities of biodiversity institutions using the R package ‘CoordinateCleaner’ v. 2.0-18 (Zizka et al., 2019).

We calculate grass species range size using an alpha hull=2 via R package ‘ConR’ v. 1.2.2 (Dauby, 2018). We used alpha hull function because it is a conservative measure that provides information on patterns of occupancy within the species ranges (Ficetola et al., 2014). As the alpha hull function is limited by occurrence data ranging within 180 degrees, widespread indigenous species whose occurrence data ranged over 180 degrees longitude (i.e., *Eleusine coracana* subsp. *africana* (Kenn.-O’Byrne) Hilu & de Wet, *Eragrostis curvula* (Schrad.) Nees, *Themeda triandra*) had their longitudinal range sizes sampled in two or three sets which were then summed together to give the final value. One set was restricted to -18.0 to 145.9 longitude (covering GBIF occurrences from e.g. the westernmost point of Africa to easternmost point of Hokkaido, Japan). Another set covered -180.0 to -35.0 longitude (covering GBIF occurrences from e.g. the Pacific and North, Central and South America). For *Eragrostis curvula* and *Themeda triandra*, where the number of cleaned GBIF occurrences was too large to compute using the R package ‘ConR’ v. 1.2.2 (Dauby, 2018), a further set was used which ranged from 146.0 to 180.0 longitude (covering GBIF occurrences from e.g. central Australia to the Fijian islands of Vanua Levu, Rabi, and Taveuni).

The mean alpha hull=2 values of all native grass species per quadrat were then calculated. Thirteen quadrats had purely alien grass species and were not included in comparisons of native grass species range sizes. To compare range sizes of just sub-Saharan endemic grasses found in the plots, widespread native grass species whose distributions extend outside sub-Saharan Africa (i.e., *Eleusine coracana* subsp. *africana*, *Eragrostis curvula*, *Lachnagrostis lachnantha, Themeda triandra*) were removed. As a result, an additional two quadrats (i.e. 15 quadrats in total) were not included in comparisons of mean sub-Saharan endemic grass species range sizes per plot.

### 2.5 Data analyses

All analyses were performed using R version 4.0.3 (R Core Team, 2020), with the individual packages used mentioned below.

#### 2.5.1 Checking correlations of environmental variables

Correlations between environmental variables recorded per plot were checked via the ‘rcorr’ function of R package ‘Hmisc’ v 4.4-1 (Harrell, 2020), by computing a matrix of Pearson’s r and Spearman’s rho (ρ) rank correlation coefficients, as well as being checked visually using R package ‘corrplot’ v 0.84 (Wei & Simko, 2017; Fig. S1). Sixteen highly correlated (p <0.05) variables were removed, i.e. ‘% plot animal disturbance’ (correlated with ‘grazing regime’), ‘elevation’ and BioClim (Table S1) variables ‘BIO.2’, ‘BIO.5’, ‘BIO.6’, ‘BIO.7’, ‘BIO.8’, ‘BIO.9’, ‘BIO.10’, ‘BIO.11’ (correlated with ‘BIO.1’), ‘BIO.14’, ‘BIO.15’, ‘BIO.17’, ‘BIO.19’ (correlated with ‘BIO.4’), BIO.16, BIO.18 (correlated with ‘BIO.13’). This left 13 remaining variables to be included in CCA: ‘burning regime’, ‘grazing regime’, ‘slope aspect’, ‘slope gradient’, ‘% bare soil’, ‘% litter’, ‘% plot burnt’, ‘% rock’, and five BioClim variables i.e. ‘BIO.1’, ‘BIO.3’, ‘BIO.4’, ‘BIO.12’, ‘BIO.13’ (Table S1).

#### 2.5.2 Examining grass species composition

To compare grass species composition of quadrats across the ecotypes and disturbance regimes, canonical correspondence analysis (CCA; Ter Braak, 1986) was performed on the data set with down-weighting of rare taxa using R package ‘vegan’ v. 2.5-6 (Oksanen et al., 2019). Grass species cover data were square-root transformed and scaled to unit variance prior to analysis. Detrended correspondence analysis (DCA) and non-metric multidimensional scaling (NMDS) were also performed but, since they gave qualitatively identical results, only the CCA results are shown here. To evaluate which environmental vectors were significant in explaining the grass species composition of plots, abiotic variables that were not highly correlated (see above) were recorded for each quadrat, and were plotted on to the CCA.

To identify grass species characteristic of each ecotype, grazing regime and burning regime, indicator species analyses were performed using the Indval method (Dufrêne & Legendre, 1997) in R package ‘labdsv’ v. 2.0-1 (Roberts, 2019). Three separate Indval analyses were performed to compare and contrast (1) ecotypes, (2) grazing regimes, and (3) burning regimes. Indval analysis was also performed on the combined ecotype-disturbance categories, but gave inconclusive results and so is not shown here. Indicator species, i.e. those with p<0.05, were categorized as to whether they were alien (according to Fish et al., 2015; Soreng et al., 2020; Sylvester et al., Accepted; see Table S2 for taxa), DMC endemic species (according to Carbutt, 2019; Linder et al., 2014; Sylvester et al., 2020; see Table S2 for taxa), and native species (mostly sub-Saharan African endemics apart from *Lachnagrostis lachnantha*, which occurs further north into Europe).

#### 2.5.3 Comparing differences in ecosystem properties between the different ecotype-disturbance categories and identifying important predictors

Normality of the variables was tested using the Shapiro-Wilk test (Mahibbur & Govindarajulu, 1997), with % cover and proportion of overall grass cover of DMC endemic grass taxa and alien grass taxa (Table S2) unable to be transformed to fit a normal distribution. Homogeneity of variance was checked using the Bartlett test and, when non-normally distributed, the Levene and Fligner-Killeen tests were used. Because all variables apart from vascular plant species richness per plot did not fit normal distribution and variance was not homogeneous, generalized linear models (GLMs) were implemented using R package ‘MASS’ v. 7.3-53 (Ripley et al., 2020), with the factor variable being ecotype-disturbance category. For vascular plant species richness data, a poisson GLM with log link function was used. For proportional data (i.e. % cover and proportion of overall grass cover of DMC endemic grass taxa and alien grass taxa) binomial GLMs were used. For mean alpha hull=2 range sizes of native and sub-Saharan African endemic grass taxa, Gamma GLMs with log link function were used. To compare differences in ecosystem properties per plot between the 22 ecotype-disturbance categories, results of GLMs were passed through two-way ANOVA with posthoc Tukey test.

To evaluate which environmental vectors were most important in explaining the patterns in each ecosystem property (i.e. vascular plant species richness, % cover and proportion of overall grass cover of DMC endemic grass taxa and alien grass taxa, mean alpha hull=2 range sizes of native and sub-Saharan African endemic grass taxa), we conducted multiple linear regressions supplemented with relative importance analysis (Grömping 2006; Tonidandel & LeBreton 2011). This was done using the matrix of environmental variables per plot that were not highly correlated (see above) as well as ecotype as a predictor. Although relative importance analysis assumes data are normally distributed, we have chosen to use them here as the analysis is fairly robust to deviations from normality. We used relative importance analysis because traditional regression cannot effectively compare the relative importance of multiple predictors. This is because correlated predictors in traditional regression analyses explain some degree of shared variance with one another. Relative importance analysis applies a transformation to each of the predictors in the linear model to make them orthogonal to each other. This allows the researcher to examine the independent amounts of variance explained by each predictor and rightfully conclude which predictors are “more important” than others within the current sample. We used the lmg method (Lindeman et al., 1980), which calculates the relative contribution of each predictor to the R square with the consideration of the sequence of predictors appearing in the model. The lmg method is considered one of the best relative importance metrics as it can decompose R^2^ into non-negative contributions that automatically sum to the total R^2^ (Grömping, 2006). We calculated the lmg and bootstrap relative importance and confidence intervals using R package ‘relaimpo’ (Grömping, 2006), with number of bootstrap replicates set at 1000 and bootstrap confidence interval type set as percentile.

#### 2.5.4 Beta diversity analyses

We chose to calculate beta diversity by comparing differences between ecotype-disturbance categories at a plot-level and at a site-level. Plot-level beta diversity more accurately represents turnover within the ecotype-disturbance categories. Site-level beta diversity more accurately represents geographic turnover of habitats between sites, although the sample size is significantly reduced. When comparing at a site-level, five of the ecotype-disturbance categories had only one observation (i.e. grassland moderate grazing low burning, shrubland moderate grazing low burning, shrubland heavy grazing high burning, wet moderate grazing high burning, wet heavy grazing moderate burning), and so could not be included in analyses, thus yielding a total sample size for analysis of 53 based on 17 ecotype-disturbance categories (see y axis of associated figure for sample size per ecotype-disturbance category used in analysis).

We calculated pairwise ecotype-disturbance category Sørensen dissimilarities (βsor) for grass species using the function beta.pair in R package ‘betapart’ v. 1.5.2 (Baselga et al., 2020). For plot-level analyses of all grass taxa, this yielded a total βsor sample size of 8316 (378 for each of the 22 ecotype-disturbance categories). For site-level analyses of all grass taxa, this yielded a total βsor sample size of 170 (10 for each of the 17 ecotype-disturbance categories). To look at what contribution turnover (βsim) and nestedness (βsne) had to overall βsor, we calculated total βsim and βsne for each ecotype-disturbance category using function beta.multi in R package ‘betapart’ v. 1.5.2 (Baselga et al., 2020).

We then compared differences in βsor of the ecotype-disturbance categories at plot- and site-level via two-way ANOVA on binomial GLM results followed by Tukey test.

As notable differences were seen in plot-level βsor across the ecotype-disturbance categories, we chose to disentangle the effects of native and exotic species on plot-level βsim and βsne by performing separate analyses for native species and alien species using the methodology described above. When comparing alien species, of the nine ecotype disturbance categories where alien species were present, three of these had only one observation (i.e. grassland low grazing moderate burning, grassland low grazing high burning, wet moderate grazing moderate burning) and so could not be included in analyses, thus yielding a total plot sample size for analysis of 38 based on six ecotype-disturbance categories (see y axis of accompanying figure for sample size per ecotype-disturbance category used in analysis). Calculating pairwise dissimilarities of these 38 plots with alien grasses yielded a total βsor sample size of 468 (78 for each of the six ecotype-disturbance categories). When comparing native species, all 22 ecotype disturbance categories were present and the total sample size for analysis was 209 (see y axis of accompanying figure for sample size per ecotype-disturbance category used in analysis). Calculating pairwise dissimilarities of these 209 plots with native grasses yielded a total βsor sample size of 8316 (378 for each of the 22 ecotype-disturbance categories).

To evaluate which environmental vectors were most important in explaining the patterns in βsor, we conducted partial Mantel tests (Smouse et al., 1986) for each environmental vector that was not highly correlated (see 2.5.1) as well as ecotype as a predictor. The dependence between distance matrices of βsor and individual environmental vectors were assessed while controlling the effect of a third geographic distance matrix. For the geographical distance matrix, paired geodesic distances between each plot (for plot-level analyses) or study site (for site-level analyses) were calculated using R package ‘geodist’ v. 0.0.7 (Padgham & Sumner, 2021). For the environmental vectors, these were converted into Euclidean distance matrices using function ‘dist’ of R package ‘vegan’ v. 2.5-6 (Oksanen et al., 2019). Partial Mantel tests were conducted using function ‘mantel’ of R package ‘ecodist’ v. 2.0.7 (Goslee & Urban, 2007), with Pearson correlation rank distances used, the number of permutations set at 10,000, and the number of iterations to use for the bootstrapped confidence limits set at 500.

## 3 Results

### 3.1 Grass species composition

In the CCA, axes 1 and 2 accounted for 22% and 19.5% of the total variance, respectively (Fig. 2). When plotting environmental vectors onto the CCA, all vectors were significant (p <0.05; Table S3), although the R^2^ values of % rock, % soil, % litter, % plot burnt were very low at < 0.1, and so were not considered as important determinants of grass community composition. When considering the other vectors, ‘BIO.1’ i.e. annual mean temperature, ‘slope’ i.e. slope gradient, ‘BIO.3’ i.e. isothermality (Table S1), and ‘grazing regime’ had the highest R^2^ values (Table S3).

**Figure 2.**
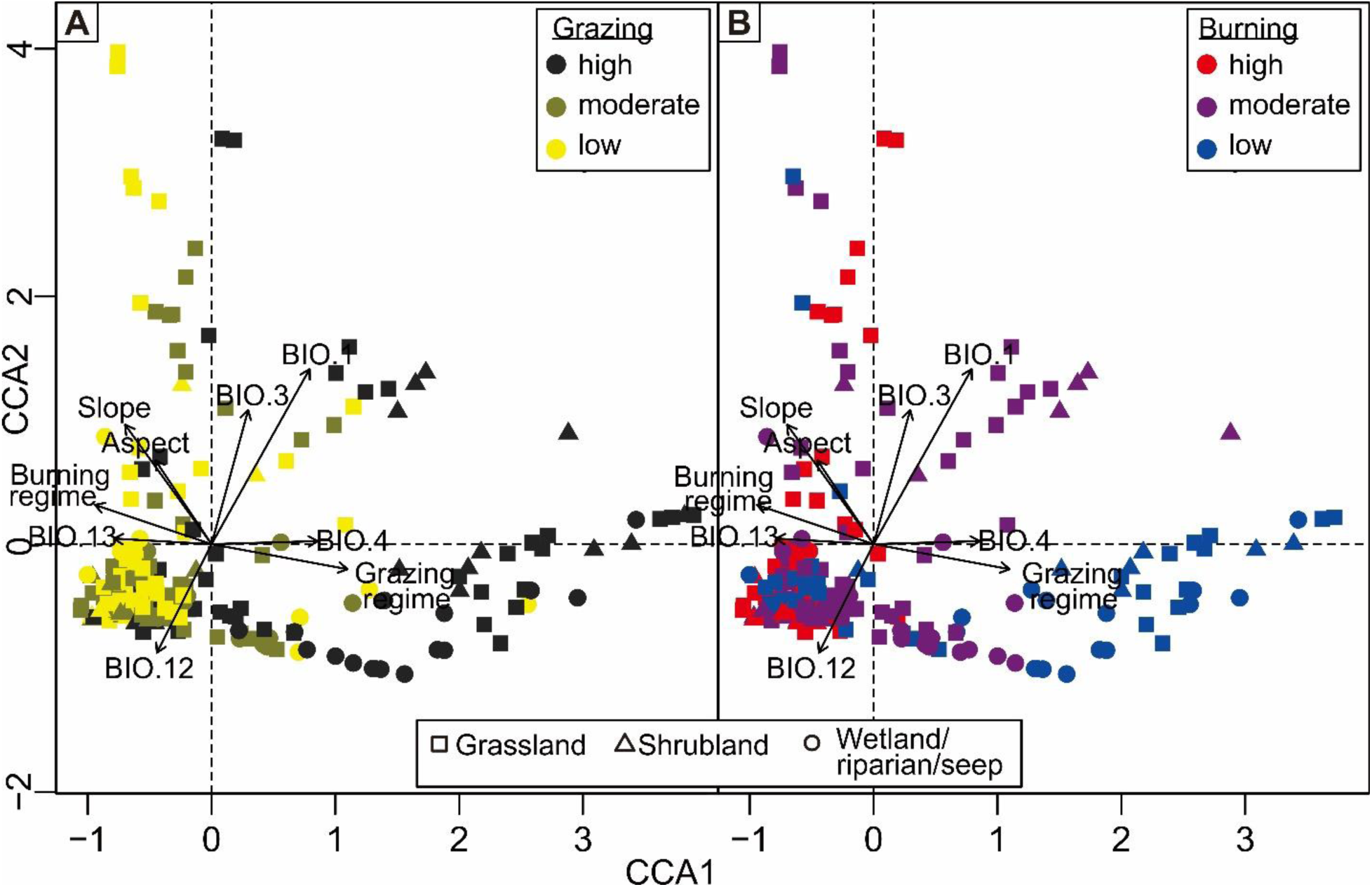
Relative positions of the study plots along axes 1 and 2 from the CCA, with shapes indicating ecotypes and colours indicating (A) grazing regimes or (B) burning regimes of the plots. Arrows refer to the environmental vectors that were significant and with R_2_ > 0.1.

A large number of mostly grassland and shrubland plots clustered on the negative axes of CCA1 and CCA2, these having mostly low and moderate (and some heavy) grazing regimes (Fig. 2A) and a mixture of burning regimes (Fig. 2B). These plots were from numerous different sites across the DMC. Environmental vectors ‘BIO.12’ (i.e. annual precipitation) and ‘BIO.13’ (i.e. precipitation of wettest month) were slightly positively associated to this large cluster, with ‘BIO.1’ (i.e. annual mean temperature) and ‘BIO.4’ (i.e. temperature seasonality; Table S1) slightly negatively associated, although no vector had a particularly large influence. One dispersed group of plots comprising a mixture of ecotypes and mostly heavy grazing regimes (Fig. 2A) and low burning regimes (Fig. 2B) was found on the positive axis of CCA1 and predominantly the negative axis of CCA2. These plots were from numerous different sites across the DMC. Environmental vectors ‘grazing regime’ and ‘BIO.4’ (i.e. temperature seasonality; Table S1) were positively associated to this group, while ‘burning regime’ and ‘BIO.13’ (i.e. precipitation of wettest month) were negatively associated. Another dispersed group of grassland and shrubland plots was found on the positive axes of CCA1 and CCA2, these being all moderately burnt (Fig. 2B) and grazing regimes changing from low to high with distance from the centre (Fig. 2A). The majority of these plots were from one site, Naudes Nek of the Eastern Cape province of South Africa. Environmental vector ‘BIO.1’ (i.e. annual mean temperature) was positively associated to this group, while ‘BIO.12’ (i.e. annual precipitation) was negatively associated. On the negative axis of CCA1 and positive axis of CCA2, a final dispersed group of mainly grassland plots can be identified which have mostly low or moderate grazing regimes and a mixture of burning regimes. The majority of these plots were from one site, Sehlabathebe National Park, Lesotho. Environmental vectors ‘slope gradient’, ‘aspect’ and ‘BIO.3’ (i.e. isothermality; Table S1) were slightly positively associated to this group, while ‘BIO.12’ (i.e. annual precipitation) was slightly negatively associated, although no vector had a particularly large influence.

When comparing grass indicator taxa (i.e. retrieved as significant, p<0.05, in indicator species analyses) between ecotypes (Fig. 3; Table S4), grassland only had one indicator, the DMC endemic *Tenaxia disticha* var. *dracomontana* H.P. Linder. Shrubland had four indicator taxa, including two natives and the alien *Poa pratensis* subsp. *pratensis* L., although the indicator taxon with the highest indval value was the DMC endemic *Tenaxia disticha* var. *lustricola* H.P. Linder. Wetland had the most indicator taxa, although none were DMC endemics, with five being native and dominating the cumulative total of indval values, and one being (a purported) alien, *Deschampsia cespitosa* (L.) P. Beauv.

**Figure 3.**
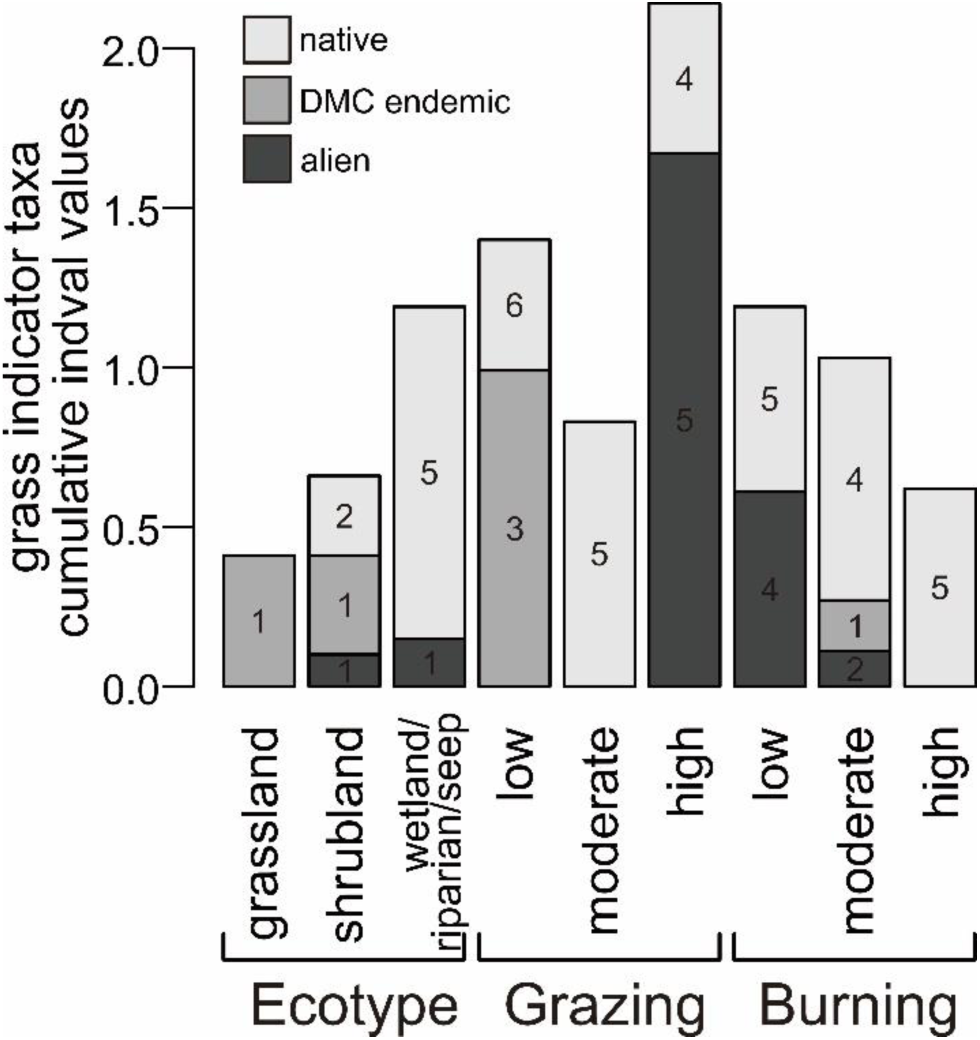
Cumulative indval values of alien (black), DMC endemic (grey), and native (white) grass indicator taxa found to be significant (p<0.05) in three different indicator species analyses comparing ecotypes, grazing regimes, and burning regimes. Numbers within the bars refer to the total number of indicator taxa pertaining to each category. Detailed information can be found in Table S3.

When comparing grass indicator taxa between grazing regimes (Fig. 3; Table S4), DMC endemic taxa were only found in low grazing, with three endemic taxa (*Festuca caprina* var. *macra* Stapf; *Tenaxia disticha* var. *dracomontana*; *Festuca caprina* var. *irrasa* Stapf) contributing a large amount to the cumulative indval value compared with the six native taxa also found as indicators of low grazing. Alien indicator taxa were only found in heavy grazing, with five alien taxa (*Agrostis gigantea* Roth; *Bromus catharticus* Vahl; *Poa annua* L.; *Poa pratensis* subsp. *pratensis*; *Poa pratensis* subsp. *irrigata* (Lindm.) H. Lindb.) contributing to a large proportion of the overall cumulative indval value when compared with the four native taxa also found as indicators of high grazing. Moderate grazing regimes had only native grass taxa as indicators.

When comparing grass indicator taxa between burning regimes (Fig. 3; Table S4), only one DMC endemic indicator taxon was found, this being the recently described *Festuca drakensbergensis* Sylvester, Soreng & M.D.P.V. Sylvester (Sylvester et al., 2020) which was an indicator of moderate burning albeit proportioning a small amount to the overall cumulative indval value. Conversely to what was found when comparing grazing regimes, alien grasses were notable in sites with low burning intensity and less so in moderate burning. There was a correlation between alien and native indicator taxa of heavy grazing and those of low burning. Seven of the nine indicator species of heavy grazing (i.e. *Agrostis subulifolia* Stapf, *Bromus catharticus*, *Catalepis gracilis* Stapf & Stent, *Eragrostis caesia* Stapf, *Poa annua*, *Poa pratensis* subsp. *irrigata*, *Tribolium purpureum* (L. f.) Verboom & H.P. Linder) were also indicator species of low burning, with the other two (i.e. *Agrostis gigantea*, *Poa pratensis* subsp. *pratensis*) being indicators of moderate burning (Table S4). In moderate burning, native indicator species contributed the most to the overall cumulative indval value, and native taxa were the only indicator species for high burning.

### 3.2 Comparison of plant species richness, grass community composition and range size between ecotype-disturbance categories and identifying important predictors

When comparing total vascular plant species richness per plot between the ecotype-disturbance categories (Fig. 4A), there were very few significant differences. However, when comparing between burning regimes within singular grazing categories, it was notable, albeit usually not significantly, that plant species richness increased with burning. This was most notable in heavily grazed grassland. Grazing did also appear to somewhat reduce plant species richness when compared within burning regimes, although this was less notable. Relative importance analyses (Fig. 5A) found numerous environmental vectors to influence plant species richness, with ‘burning regime’, followed by ‘% rock’ and ‘slope gradient’, the most important.

**Figure 4.**
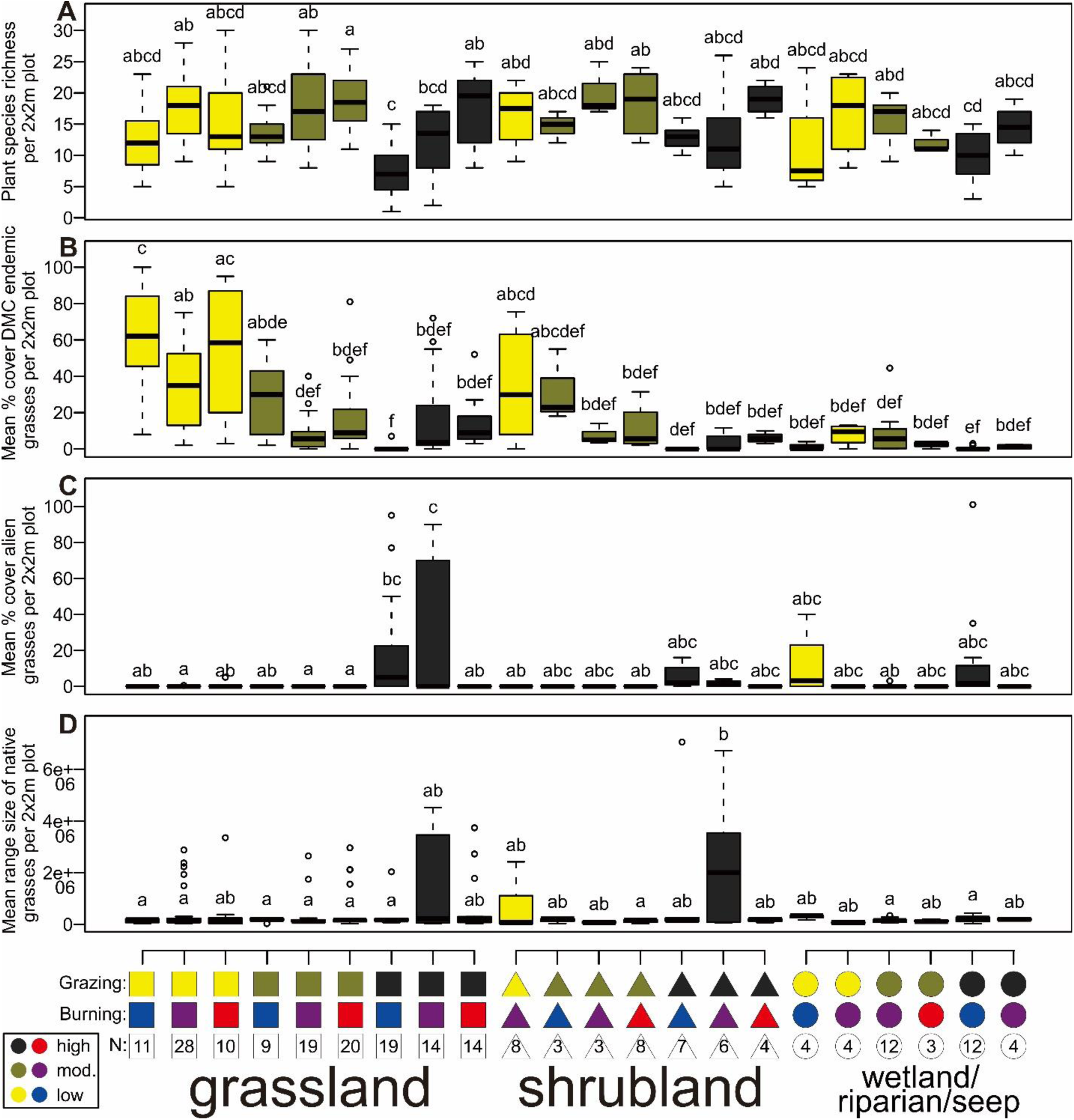
Plant diversity, composition and range size traits across different ecotype and disturbance regimes (A-C). Comparisons of (A) total vascular plant species richness per plot; (B) % cover of grass taxa endemic to the DMC per plot; (C) % cover of alien grass taxa per plot (see Table S2 for taxa); (D) mean range size, calculated as alpha hull=2, of native grass taxa per plot. Boxplots are colour coded according to grazing regime, with colour codes of grazing and burning regimes found on the x axis as well as number of 2×2m plots (N) also indicated. Different letters above boxplots denote significant (p <0.05) differences following two-way ANOVA on GLM results and post-hoc Tukey test.

**Figure 5.**
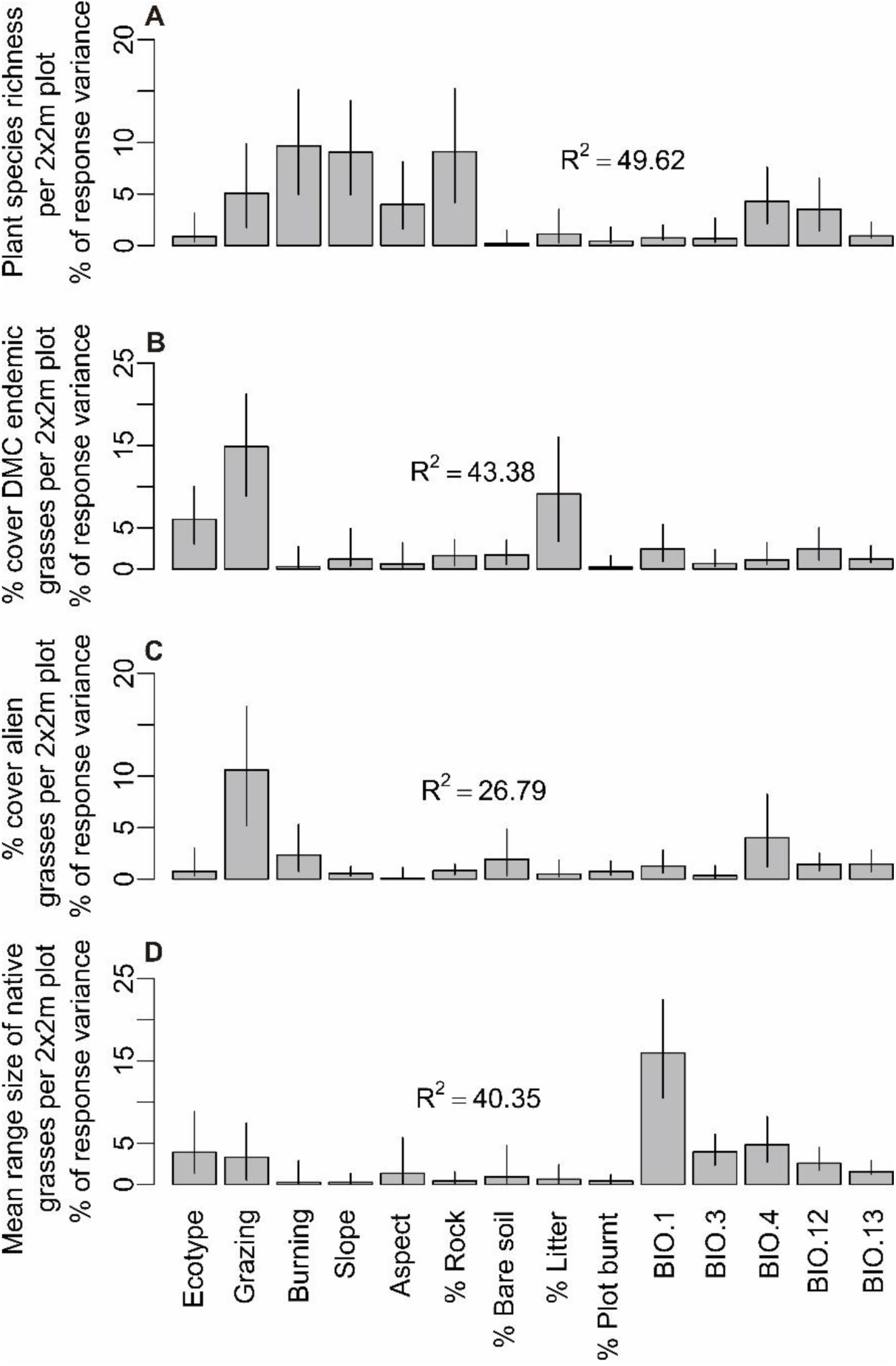
Relative importance of environmental vectors on plant diversity, composition and range size traits (A-D). Shows % response of variance and 95% confidence intervals of environmental vectors when analysing (A) total vascular plant species richness per plot; (B) % cover of grass taxa endemic to the DMC per plot; (C) % cover of alien grass taxa per plot (see Table S2 for taxa); (D) mean range size, calculated as alpha hull=2, of native grass taxa per plot. Values based on relative importance analysis using the lmg method. R^2^ values of each analysis are shown within plots. Description of BioClim variable codes found in Table S1.

When comparing mean % cover of DMC endemic grasses per plot between the ecotype-disturbance categories (Fig. 4B), there was a difference between grazing regimes, most notably that of grassland and shrubland, with greater cover of DMC endemics being found in sites with low grazing. However, statistical support for this was generally low, with only a few differences being significant, e.g. low-grazed low-burnt grassland differed from all other grassland disturbance categories apart from low-grazed high-burnt grassland. Looking at the proportion of grasses which are DMC endemics across the ecotype-disturbance categories (Fig. S2A), the differences are less apparent, although the above-mentioned patterns are still somewhat evident. Relative importance analyses (Fig. 5B) found ‘grazing regime’, followed by ‘% litter’ and ‘ecotype’, to be the most important environmental vectors explaining the patterns in mean % cover of DMC endemic grasses per plot. When looking at proportion of grasses which are DMC endemics per plot, relative importance analyses found a similar pattern to the above, with grazing regime having the highest percentage of response variance, although this was followed by ‘ecotype’ and then ‘BIO.1’, with ‘% litter’ of little importance, but other vectors having a higher percentage of response variance (Fig. S3A).

When comparing mean % cover of alien grasses per plot (Fig. 4C) and the proportion of grasses which are alien per plot (Fig. S2B) between the ecotype-disturbance categories, aside from low-grazed and low-burnt wet ecotypes, alien grasses were only notable in heavily grazed sites with low or moderate burning of each ecotype. The only significantly different categories were heavily-grazed and low- or moderately-burnt grasslands when comparing mean % cover of alien grasses per plot (Fig. 4C). However, when comparing the proportion of grasses which are alien per plot (Fig. S2B), heavy-grazed low-burnt shrubland and wet ecotypes were also retrieved as significantly different. Relative importance analyses (Fig. 5C) found ‘grazing regime’ to be the most important environmental vector explaining the patterns in mean % cover of alien grasses per plot, with this followed by ‘BIO.4’, ‘burning regime’, and ‘% bare soil’. When looking at proportion of grasses which are alien per plot, relative importance analyses found ‘grazing regime’ followed by ‘burning regime’ to be the most important vectors in explaining the variance in the dataset (Fig. S3B).

When comparing mean range size of native grasses per plot between the ecotype-disturbance categories (Fig. 4D), shrubland with heavy grazing and moderate burning was significantly different from the other categories. Grassland with heavy grazing and moderate burning, and to a lesser extent shrubland with low grazing and moderate burning, also showed a greater mean range size per plot, although this was not statistically significant. When only comparing mean range sizes of sub-Saharan endemics, with the widespread native taxa (i.e., *Eleusine coracana* subsp. *africana*, *Eragrostis curvula*, *Lachnagrostis lachnantha, Themeda triandra*) removed, no significant differences were observed between the ecotype-disturbance categories (Fig. S2C). Relative importance analyses (Fig. 5D, S3C) found ‘BIO.1’ to be disproportionately larger than the other environmental vectors in its percentage of response variance in explaining the patterns when comparing mean range size of native and just sub-Saharan endemic grasses per plot between the ecotype-disturbance categories.

### 3.3 Beta diversity

Comparing βsor of all grasses at the plot-level across the ecotype-disturbance categories (Fig. 6A), heavily-grazed grassland, shrubland, and partly wet habitat, plots often had the highest βsor. This pattern was more erratic in shrubland and wet habitat ecotypes, with ecotype disturbance categories with low sample size, i.e. three or four plots, generally having lower βsor values. βsim (i.e. turnover) generally contributed the most to overall βsor at the plot-level (Fig. 6B), with βsne (i.e. nestedness) only forming an important component in ecotype-disturbance categories with low overall βsor, these with low sample sizes three or four plots. Partial Mantel tests found ‘BIO.1’, followed by ‘grazing regime’, to be the most important in explaining plot-level βsor of all grasses, although other vectors were also significant, i.e. (in order of highest to lowest Mantel coefficient values) ‘% bare soil’, ‘ecotype’, ‘burning regime’, ‘aspect’, ‘BIO.13’, ‘BIO.12’, ‘BIO.4’ (Table 1).

**Figure 6.**
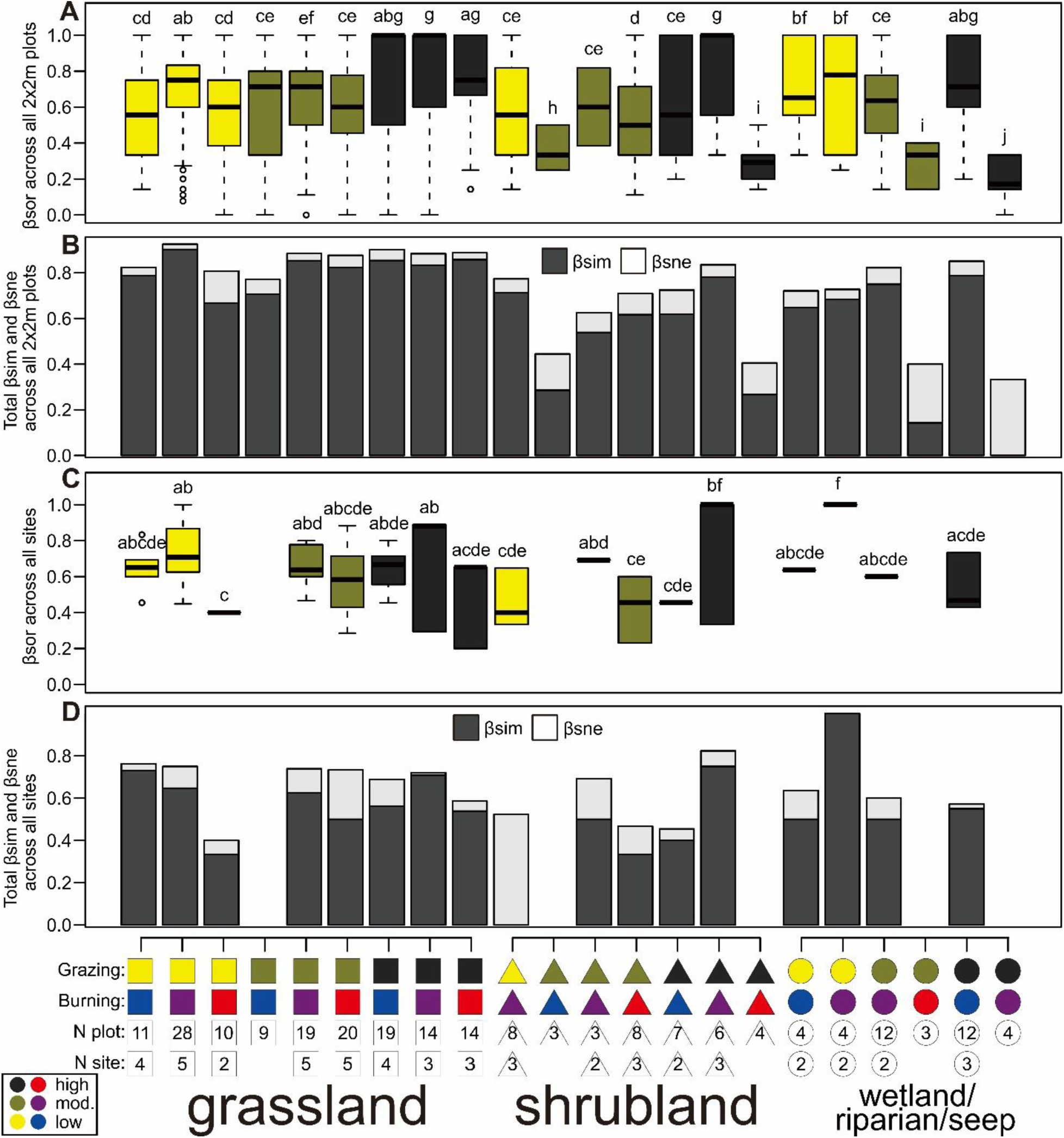
Grass community Beta diversity traits across different ecotype and disturbance regimes at a plot-level (A-B)and site-level (C-D). Comparisons of (A) plot-level beta diversity of grass taxa, represented by overall Sørensen dissimilarity (βsor) of ecotype-disturbance categories across 2×2m plots; (B) contribution of turnover (βsim) and nestedness (βsne) to overall plot-level βsor; (C) site-level beta diversity of grass taxa, represented by overall βsor of ecotype-disturbance categories across study sites; (D) contribution of βsim and βsne to overall site-level βsor. Boxplots of (A) and (C) are colour coded according to grazing regime, with colour codes of grazing and burning regimes found on the x axis as well as sample size i.e. number of 2×2m plots (N plot) or number of sites (N site). Different letters above boxplots of (A) and (C) denote significant (p <0.05) differences following two-way ANOVA on binomial GLM results and post-hoc Tukey test.

**Table 1.** Mantel coefficients showing the effects of individual environmental vectors, given geographic distance, on βsor for all, native and alien grasses at plot-level and all grasses at site-level. Significant (two-tailed p <0.05) Mantel coefficients are in bold. Signif. codes: ‘***’=p<0.001, ‘**’=p<0.01,’*’=p<0.05, ‘.’=p<0.1.

| | βsor<br>plot-level all<br>spp.<br>(N=222) | βsor<br>plot-level<br>natives<br>(N=209) | βsor<br>plot-level<br>aliens<br>(N=41) | βsor<br>site-level<br>all spp.<br>(N=53) |
| --- | --- | --- | --- | --- |
| Ecotype | **0.14\*\*\*** | **0.16\*\*\*** | **0.11\*** | **0.19\*\*** |
| Grazing regime | **0.17\*\*\*** | **0.16\*\*\*** | **0.27\*\*\*** | **0.16\*\*\*** |
| Burning regime | **0.14\*\*\*** | **0.12\*\*\*** | **0.52\*\*\*** | **0.20\*\*\*** |
| Slope gradient | 0.04 | **0.07\*** | **0.12\*** | **0.17\*** |
| Aspect | **0.11\*\*\*** | **0.13\*\*\*** | 0.03 | **0.19\*\*** |
| % Rock | -0.02 | -0.01 | **0.15\*** | 0.04 |
| % Bare soil | **0.16\*\*\*** | **0.12\*\*** | **0.16\*** | **0.24\*\*** |
| % Litter | -0.03 | -0.03 | **0.35\*\*\*** | 0.07 |
| % Plot burnt | -0.05 | -0.04 | 0.09 | -0.08 |
| BIO.1 | **0.21\*\*\*** | **0.22\*\*\*** | 0.05 | **0.19\*\*** |
| BIO.3 | -0.04 | -0.04 | **-0.17\*** | -0.07 |
| BIO.4 | **0.06\*** | 0.02 | **0.20\*\*** | 0.11 |
| BIO.12 | **0.09\*\*\*** | **0.13\*\*\*** | -0.05 | -0.03 |
| BIO.13 | **0.11\*\*\*** | **0.12\*\*\*** | **0.11\*** | 0.03 |

When looking at βsor of native and alien grasses at the plot-level across the ecotype-disturbance categories (Fig. S4), the strong patterns of notably higher βsor in heavily grazed ecotypes were not so evident. Nevertheless, plots of heavily-grazed low burnt grassland, shrubland, and wet habitat had the highest βsor when considering just native grasses (Fig. S4C) despite omission of alien grasses which also exhibited the high βsor values for these ecotype-disturbance categories (Fig. S4A). βsim generally contributed the most to overall βsor of native grasses at the plot-level (Fig. S4D), with βsne only forming an important component in shrubland and wetland disturbance categories with low overall βsor, these with low sample sizes three or four plots. For alien grasses, βsim contributed the most to overall βsor in heavily grazed grassland and wet habitat ecotypes, although βsne formed an important component in heavily grazed shrubland, these with low overall βsor and low sample sizes four or five plots (Fig. S4B). Partial Mantel tests on plot-level βsor of native grasses found similar or more pronounced influence of environmental vectors compared with that on plot-level βsor of all species, with ‘BIO.1’, followed by ‘grazing regime’ and ‘ecotype’, being the most important, although ‘BIO.4’ no longer had an influence (Table 1). When considering plot-level βsor of alien grasses, partial Mantel tests found ‘burning regime’ to have a disproportionately higher influence, followed by ‘% litter’, ‘grazing regime’, and ‘BIO.4’ (Table 1). Other vectors were also significant, i.e. (in order of highest to lowest Mantel coefficient values) ‘BIO.3’, ‘% bare soil’, ‘% rock’, ‘slope gradient’, ‘ecotype’ (Table 1). Certain vectors found to be significant for all and native species at plot-level e.g. ‘BIO.1’, ‘BIO.12’ and ‘slope aspect’, were not significant for plot-level βsor of alien grasses (Table 1).

Comparing βsor at the site-level across the ecotype-disturbance categories (Fig. 6C), no notable differences were seen within the grassland ecotype, which was the most heavily sampled at site-level, apart from low-grazed high-burnt grassland, which had a significantly lower βsor but this also with the lowest sample size (N=2) among the grassland disturbance categories. Shrubland disturbance categories also did not differ from those of the majority of grassland apart from heavy-grazed moderately-burnt shrubland, which was significantly higher than the other shrubland disturbance categories, and many of the grassland disturbance categories as well as heavy-grazed low-burnt wet habitat. Wet habitat disturbance categories also did not differ from those of the majority of grassland and shrubland apart from low-grazed moderate-burnt wet habitat, which was significantly higher and with 100% turnover (Fig. 6D). Similar to plot-level analyses, βsim generally contributed the most to overall βsor at the site-level (Fig. 6D). βsne only dominated in low-grazed moderately-burnt shrubland. Partial Mantel tests on site-level βsor of all grasses found ‘% bare soil’ to have the greatest influence, followed by ‘burning regime’, ‘ecotype’, ‘aspect’, ‘BIO.1’, ‘slope gradient’ and ‘grazing regime’ (Table 1). Apart from ‘BIO.1’, no other bioclimatic variables were retrieved as significant.

## 4 Discussion

### 4.1 Notable detrimental impact of heavy grazing on DMC Afroalpine vegetation

Our comparisons of plant/grass diversity and composition found little differences between most ecotype-disturbance categories, with only heavy-grazing regimes having a notable impact, and that mainly on the grassland ecotype. It must be understood, however, that heavy-grazing regimes and the grassland ecotype are the most widespread and prevalent across the DMC Afroalpine landscape (Carbutt, 2020; Turpie et al., 2021). Thus, landscape-scale extrapolations will likely show an immense anthropogenic impact, although we have not undertaken spatial analysis of this.

We show here that heavily grazed grass communities exhibit a distinct composition in CCA (Fig. 2): one notable groups of plots separated from the large group of plots, that represent the more uniform Afroalpine grass community, and were positively associated to grazing regime and BIO.4 (i.e. temperature seasonality) and negatively associated to burning regime and BIO.13 (i.e. precipitation of wettest month). These plots had mostly high-grazing and low-burning regimes, with these corresponding to areas so heavily grazed that there was insufficient fuel load to support fires and are here referred to as ‘short grazing lawns’. The relationship between high-grazing and low-burning regimes seen in the CCA (Fig. 2) is also notable by how seven out of nine indicator species of high grazing were also indicators of low burning (Fig. 3; Table S3), with the majority of these being alien.

The detrimental impact of intense grazing is shown by how alien grasses (Fig. 4C, S2B), and in isolated cases widespread natives (Fig. 4D), significantly increase, while DMC endemic grasses are reduced (Fig. 4B, S2A) in heavily grazed areas, particularly those of grasslands. Likewise, the prominent indicator species of heavily grazed areas were alien, while low grazed areas hosted DMC endemics (Fig. 3). Landscape scale extrapolations of this will likely show human impact through grazing leads to large-scale shifts in grassland communities favouring alien and widespread generalist species to the detriment of rare and endemic species, as has been shown in the Peruvian Andes (Sylvester et al., 2017). Ecotypes that were exceptionally grazed such that there was insufficient fuel load for burning (i.e. most heavy-grazed low-burnt grassland and wetland plots) also often had the lowest plant species diversity (Fig. 4A). This would imply that the vast grazing lawns present in much of the DMC Afroalpine landscape possibly have significantly lower diversity compared to less-grazed areas and that intense grazing may lower diversity at a landscape scale. Negative impacts of intense grazing on plant diversity, particularly that of forbs, have also been found in studies from lower-elevation grasslands in South Africa (reviewed in O’Connor et al., 2010).

### 4.2 Resilience of DMC Afroalpine plant diversity and composition to moderate grazing and intense burning

Human impact through grazing and burning is often considered a primary influencing factor on the biodiversity of high elevation ecosystems (Körner 2003). Indeed, heavy-grazing regimes had a notable detrimental impact mainly on the grassland ecotype, with the landscape-scale consequences of this possibly immense due to the ubiquity of heavily-grazed grasslands in the DMC, as mentioned in the previous section. Nevertheless, there was also a notable lack of significant differences between many plant diversity and composition traits, and in some cases there even being a positive influence of grazing or burning, that does imply the DMC Afroalpine vegetation is fairly resilient to grazing and burning.

This notable lack of significant differences between most ecotype-disturbance categories can be seen in plot comparisons of vascular plant species richness, average range sizes of native and sub-Saharan grasses and site-level βsor across all ecotypes, and cover and proportion of DMC endemic and alien grasses per plot of shrubland and wet habitat ecotypes (Fig. 4, 6C, S2). CCA also found a large group of mainly grassland and shrubland plots of mainly low- and moderate-grazing and differing burning regimes with no particular environmental vector associated to them (Fig. 2) and which could be seen to represent a more uniform Afroalpine grass community. Two other groups of plots were also differentiated based on environmental vectors not related to grazing and burning (Fig. 2). This influence of other factors besides grazing and burning on plant diversity and composition can also be seen by how numerous other environmental vectors were found to be the principal determinants of plant species richness and mean range size of native and sub-Saharan endemic grasses per plot in relative importance analyses (Fig. 5A, D, S3C) and plot- and site-level βsor in partial Mantel tests (Table 1). Furthermore, cover and proportion of DMC endemic and alien grasses was also influenced by other vectors besides grazing and burning (Fig. 5B-C, S3A-B). Most notably this includes ecotype, litter cover, BIO.1 (i.e. annual mean temperature), BIO.12 (i.e. annual precipitation) and slope gradient and aspect for DMC endemics (Fig. 5B, S3A), and BIO.4 (i.e. temperature seasonality), cover of bare soil and rock, and BIO.1. Other vectors that exhibited high collinearity with BIO.1, BIO.4 and BIO.13 (Fig. S1) and, thus, were removed from analyses (see section 2.5.1) should also be borne in mind. This all indicates that diversity in these landscapes is driven by environmental heterogeneity (Tamme et al., 2010; Stein et al., 2014).

In some instances, human-induced disturbance through grazing and burning even had a positive impact on DMC Afroalpine plant diversity. This can be seen by plant species richness increasing with burning under certain ecotype-grazing categories, most notably that of heavily grazed grassland (Fig. 4A), and plot-level βsor of all grasses and native grasses increasing under heavy grazing, particularly for grassland (Fig. 6A, S4C). Other studies in Afroalpine vegetation have also found human-induced burning to have a positive effect on diversity by maintaining a structurally and biologically diverse environment (Wesche et al., 2000; Johansson et al., 2012, 2018). Human-induced burning has even been noted to increase the available habitat for less competitive high-elevation plants, thus maintaining the alpine niche or extending it to lower elevations and consequently buffering threats from climate change (Johansson et al. 2018). Burning also speeds below-ground nutrient cycling that favours the health of fire-tolerant plants (Binkley & Fisher, 2019). However, little is known regarding what would be the optimal burning regime for balancing grazing and biodiversity within DMC Afroalpine vegetation. Our data seems to suggest that annual burning regimes promote the highest diversity, but that grazing has a much stronger overriding influence on plant diversity and composition that may obscure these patterns.

For our β diversity analyses, caution must be given to comparisons of ecotype-disturbance categories with a low sample size, as these are likely to not reflect what is happening in nature, with these often having much lower βsor values at plot-level (Fig. 6A) and largely differing βsor values at site-level (Fig. 6C). Keeping this in mind and focussing on well-sampled ecotype-disturbance categories, certain patterns are still evident. Our data on plot-level βsor of all grasses (Fig. 6A), native grasses (Fig. S4C) and alien grasses (Fig. S4A), which was highest under heavy grazing regimes and with turnover usually the dominating component (Fig. 6B, S4B, S4D), shows that intense grazing leads to largely different grass species assemblages across the DMC landscape. This high plot-level βsor was particularly notable for the ‘short grazing lawns’ i.e. heavy-grazed low-burnt ecotypes where there was insufficient fuel load for burning. The highly variable composition of these ‘short grazing lawns’ was also notable in CCA (Fig. 2), where a group of heavily-grazed and low-burnt plots was highly dispersed along the CCA1 axis, indicating highly variable grass community composition.

This high variability and turnover in grass species composition across heavily-grazed low-burnt ‘short grazing lawn’ ecotypes is difficult to explain, with it also notable that these plots, especially those of grassland, often had the lowest species richness of all ecotype-disturbance categories (Fig. 4A). CCA found BIO.4 (i.e. temperature seasonality, and other vectors that were correlated with it but not included in analyses, Fig. S1; see section 2.5.1) had a strong association to the dispersion of the heavily-grazed low-burnt plots (Fig. 2; Table S3). BIO.4 is also an important vector explaining variance in the dataset for cover, proportion (Fig. 5C, S3B) and plot-level βsor (Table 1) of alien species, which often dominated heavily-grazed low-burnt plots (Fig. 3, 4C, S2B). Thus, alien grass species, and the environmental vectors associated with them, had some influence on overall plot-level βsor. Nevertheless, plot-level βsor of native taxa still had the highest values in heavily-grazed low-burnt ‘short grazing lawn’ plots (Fig. S4C). This is in contrast to the idea that human disturbance leads to biotic homogenisation (McKinney & Lockwood, 1999), with other studies finding disturbance correlates with reduced βsor (see review in Olden et al., 2018).

Heavy grazing disturbance is known to open up niches allowing numerous species to establish (Borer et al., 2014). Thus, it is more likely that differing plant communities could develop across a heavily grazed landscape, with factors determining this including local species pool, microhabitat characteristics etc. Partial Mantel tests gave the principal causal factor of high plot-level βsor in all and native grasses as BIO.1 (i.e. mean annual temperature; as well as other temperature related variables and elevation that were correlated with it, Fig. S1; see section 2.5.1), as well as numerous other vectors including grazing and burning regime (Table 1). BIO.1 was also the only bioclimatic variable that was retrieved as significant in partial Mantel tests on site-level βsor of all grasses (Table 1), which more accurately represents geographic turnover of habitats between sites. This highlights that environmental heterogeneity plays a large part, and this links with grazing and burning disturbance, to increase beta diversity in these landscapes.

It could be posited that ecosystems with a long evolutionary history of grazing and burning have vegetation which has evolved adaptations to these disturbances and, thus, can be considered more resilient. This is supported by evidence that large- or medium-sized grazing ungulates have been present in the DMC Afroalpine environment since at least ∼21,000 years ago (Grab & Nash, in press), and that natural fires were fairly common in these landscapes and ignited by lightning strikes during transition from the dry to the wet season (Archibald et al., 2012; Finch et al., in press). Pre-adaptations to mechanical disturbance by the hooves of grazing livestock may also be posited due to the mass wasting nature of montane and sub-alpine areas of the DMC that are found in precipitous mountains, while the Afroalpine DMC vegetation may be pre-adapted due to past (e.g. Last Glacial Maximum) and, more limited, current cryogenic activity (Grab et al., 2021). This is partly supported by the several annual non-grass endemics being found in the DMC, although these endemics are not a large component of the flora (Carbutt, 2019).

### 4.3 Support for the natural grass-dominated nature of the DMC

The high levels of endemism found in low or moderately disturbed grassland and shrubland (Fig. 4B, S2A), with DMC endemic indicator species mainly associated to low grazing (Fig. 3), also supports the notion that Afroalpine grassland is a natural component of the DMC (Meadows & Linder, 1993). This notion is further supported by statistically insignificant differences between ecotype-disturbance categories with regards total vascular plant species richness (Fig. 4A), native grass range sizes (Fig. 4D, S2C), site-level β diversity (Fig. 6C), and the grass community composition of the majority of plots being similar despite differing grazing and burning regimes (Fig. 2). Because of the dominance of montane forest and forest-related impacts in the origin of African mountain phytogeographical research (eastern Africa), there is the common perception that alti-montane grasslands throughout Africa are a secondary habitat following human-mediated forest removal and increased fire frequency. This is reinforced by Eurocentric thinking, stemming from the classic vegetation zones (strongly featuring forest) apparent in the Alps as being normative. Although this is certainly true and well-supported for many mountain regions globally (Körner 2003, 2012), there is a compelling body of palaeo-ecological, biogeographical and base-line biodiversity evidence showing the deep age of southern Africa’s alti-montane grasslands (Meadows & Linder, 1993; Lodder et al., 2018; Breman et al., 2019), with our research providing further support.

It must be borne in mind that our study has not included sites which could be considered truly pristine, as human impact through livestock grazing and burning is prevalent throughout the whole of the DMC. Our categorization of disturbance regimes was based on a relative scale, although it is possible that our low-grazed low-burnt category emulates pristine conditions. Nevertheless, evidence points to disturbance through grazing and burning being a natural component of these ecosystems. Paleoecological research has found that large herbivores were a common component of the DMC alti-montane grasslands (Grab & Nash, in press), although densities of these are unknown. Likewise, high-intensity late-season lightning fires are considered to be the natural fire regime in these habitats (Finch et al., in press).

### 4.4 Considerations and limitations within the dataset

Aside from other limitations and considerations already mentioned, it must be borne in mind that the responses of variables based on grass species communities to grazing and burning that we have found in this study may not reflect overall ecosystem level responses. Recent research in South African lowland savannas (Van Coller et al., 2018) and montane and coastal areas (Zaloumis & Bond, 2016) found that herbaceous responses to grazing and burning regimes differ between grasses and forbs. Nevertheless, as grasses functionally dominate the DMC Afroalpine ecosystems, the information that we provide here should not be ignored.

Care must also be taken when considering the taxonomic level of diversity that we have discussed here. While we have treated total vascular plant species richness per plot at a species-rank, we evaluate general grass community composition, cover and proportion of DMC endemic and alien grasses, range sizes and beta diversity at the variety-rank (where applicable). This is especially pertinent with regards classification of DMC endemic grasses (Table S2). Aside from *Catabrosa drakensbergensis* (Hedberg & I. Hedberg) Soreng & Fish and *Festuca drakensbergensis*, all other DMC endemics in this study are varieties of *Festuca caprina* Nees (Sylvester et al., 2020) or *Tenaxia disticha* (Nees) N.P.Barker & H.P.Linder (Linder et al., 2014). We have chosen to recognize at the varietal level as these varieties may need to be raised to the level of subspecies or species in the case of *F. caprina* var. *irrasa* and *F. caprina* var. *macra* (Sylvester et al., 2020), and autecological evidence points to the probability of the same being done for *Tenaxia disticha* var. *dracomontana* and *Tenaxia disticha* var. *lustricola* (S.P. Sylvester, pers. observation; DNA analyses of our herbarium and silica dried vouchers are planned to examine this possibility). Nevertheless, if these were treated at species level then there would be little difference in the cover and proportion of DMC endemic grasses across the ecotype-disturbance categories. Furthermore, while we have compared the known endemic taxa of the DMC (i.e. Carbutt, 2019, as well as varietal endemics and new species discovered by Linder et al., 2014, and Sylvester et al., 2020), this also needs reappraisal. Previous research that has categorized southern African grasses based on their rarity (Barker & Fish, 2007) could not be used in this study as discrepancies were present that need in-depth re-evaluation based on up-to-date distribution data and ecological knowledge, and should be the topic of future research.

Finally, it is also important to note that our fieldwork was conducted during a particularly wet year. This may have facilitated the establishment of the short grazing lawns that were notable in heavily grazed areas, especially as these tended to be dominated by annual, and often alien, grasses. In a drought year, cover of annuals would likely be minimal, exposing more ground to erosion, and driving livestock to graze on perennial grasses, with grazing impact likely to be less notable among ecotypes. Moreover, annuals only cover the ground for a portion of the year even in a good year and their root mass is comparably very low. Focused research comparing life histories and other functional traits of the grasses per ecotype-disturbance category would further improve our functional understanding of DMC Afroalpine ecosystems, and will be addressed in a subsequent paper.

### 4.5 Conclusions

Our findings of resilience of many plant diversity and composition properties to human-induced grazing and burning does provide some encouragement for conservation work in the DMC that attempts to achieve a compromise between the needs of the pastoralist community and that of biodiversity. This resilience, coupled with high levels of endemism found in less disturbed habitats, also provides support for the grass-dominated Afroalpine landscape as a natural component of the DMC, thus challenging the current narrative that tree-poor landscapes are ‘bad’ and ‘degraded’ and advising against any proposals for forestation (Zaloumis & Bond, 2016). However, the stark negative impact of heavy grazing on grassland diversity and composition, coupled with the disproportionately large area of the DMC currently occupied by heavily-grazed grassland (Carbutt, 2020; Turpie et al., 2021), is a major cause for concern.

Our results suggest that poorly-managed systems of grazing result in pressures that favour degeneration of plant communities and increases in weedy species, with this beginning to occur once a certain threshold of disturbance is passed. This ‘tipping point’ after which Afroalpine ecosystems begin to rapidly degrade could not be identified in this study as our categories for grazing and burning regimes were too broad and somewhat subjective. Comprehensive and fine-scale objective quantification of grazing and burning across different ecotypes would help shed light on these plausible ‘tipping points’ and provide scientifically sound advice to support land management strategies that mitigate against this potential ecosystem collapse. As humans are deeply ingrained in these landscapes, the more important question, it seems, is what kind of plant diversity is healthy from a socio-ecological point-of-view. Combined analyses, focusing on taxonomic and functional diversity and ecosystem services at multiple scales, would allow cost-benefit trade-offs to be quantified which would enable a more effective conservation management strategy to help safeguard this unique and fascinating vegetation-type.

## Acknowledgements

We wish to gratefully thank Nanjing Forestry University, China, and the Afromontane Research Unit (University of the Free State, South Africa) for financial and logistical support; Lyn Fish, and PRE staff for access to the PRE herbarium and discussions of taxa; and Ralph and Nadine Clark for providing an operations base in South Africa (including during lockdown). We also wish to extend grateful thanks to the permitting authorities and landowners for the relevant permits and permissions to undertake the fieldwork: Ezemvelo KZN Wildlife (uKhahlamba-Drakensberg Park & UNESCO World Heritage Site), Eastern Cape Parks & Tourism Authority, Eastern Cape Department of Economic Development, Environmental Affairs & Tourism, the Kingdom of Lesotho Department of Environment, and Witsieshoek Mountain Lodge/Batlokwa Tribal Authority.

## Author contributions

SPS, VRC and RJS conceived the ideas and designed the methodology; SPS, RJS, MDPVS and AM collected the field data, with ACM contributing native grass range size data; SPS, RJS and ACM conducted taxa identifications; SPS conducted most data analysis, with ACM obtaining alpha-hull=2 values for native grass taxa; SPS led the writing of the manuscript, with notable inputs from VRC, RJS, ACM, AM and MDPVS. All authors contributed critically to the drafts and gave final approval for publication.

## Data availability

We intend to archive our two datasets on (1) grass species composition per plot; and (2) vascular plant species richness, cover and proportion of DMC endemic and alien grasses, mean alpha hull=2 range sizes of native grasses and environmental vectors per plot in the Dryad Digital Repository.

## Supplementary Materials

**Figure S1.**
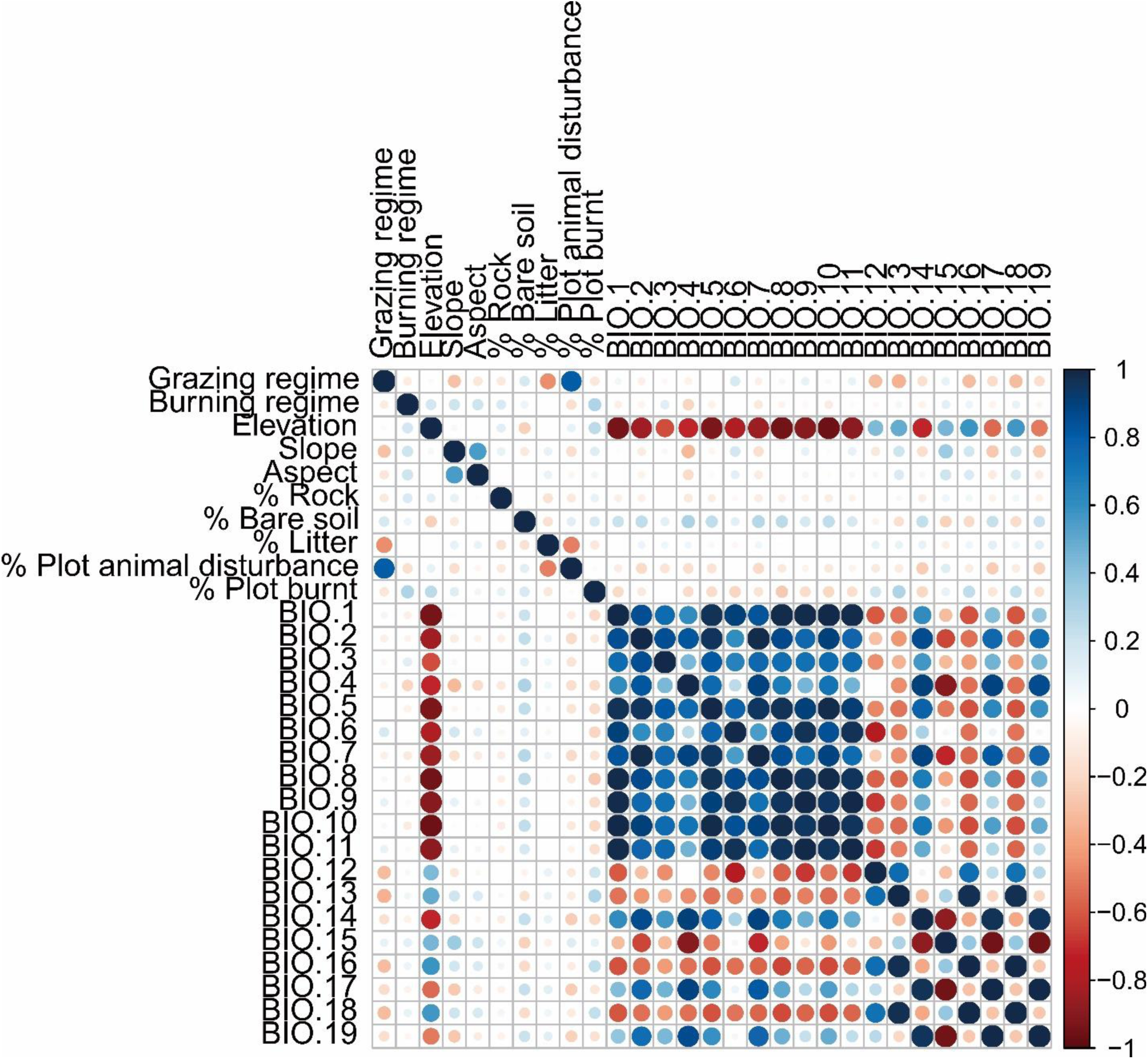
Correlogram of environmental variables based on Pearson correlation coefficients. The areas of circles show the absolute value of the Pearson correlation coefficient. Shading of circles indicates the strength of the relationship as well as the direction of the relationship i.e. positive correlations are displayed in a blue scale while negative correlations are displayed in a red scale.

**Figure S2.**
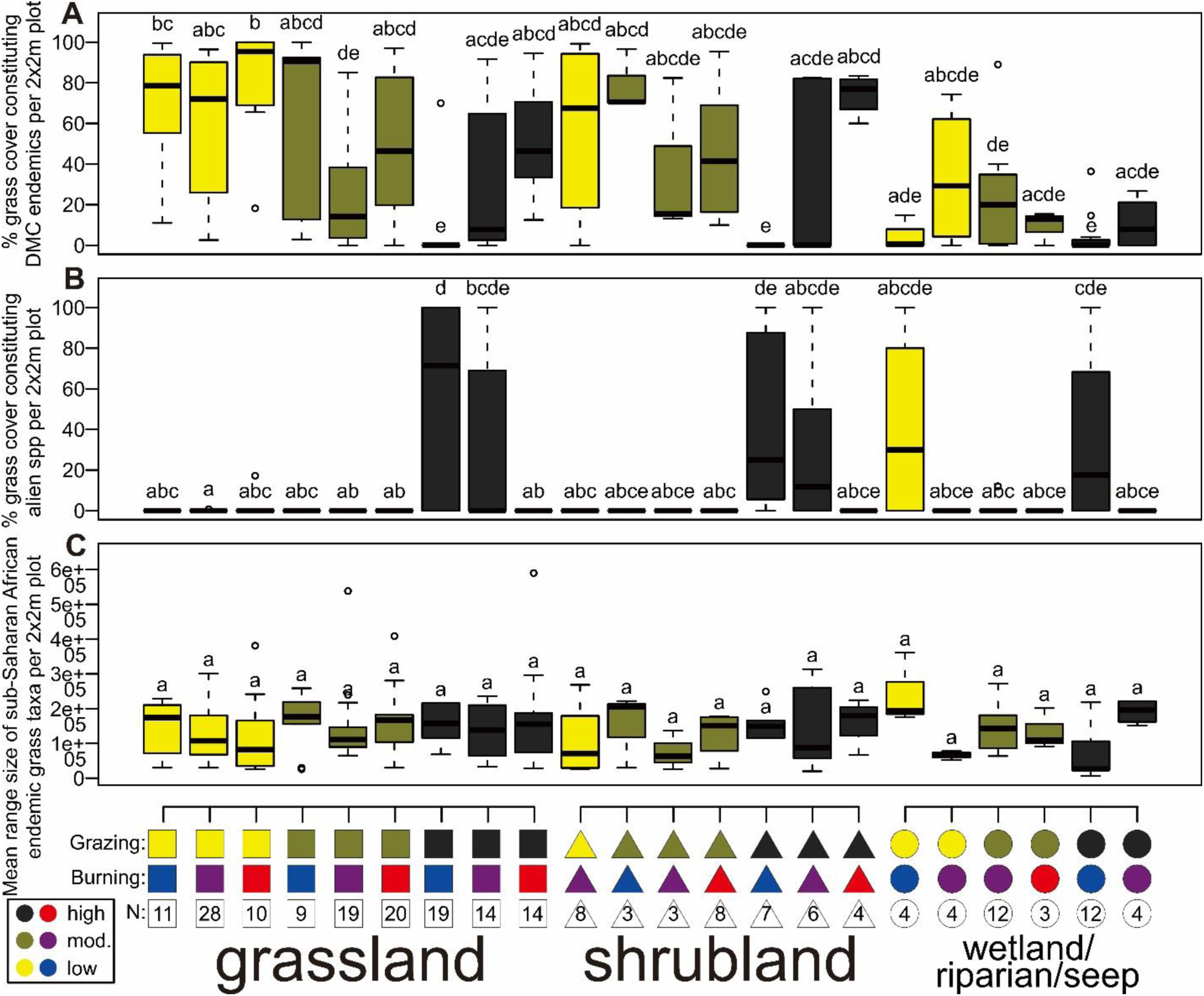
Grass community composition and range size traits across different ecotype and disturbance regimes (A-C). Comparisons of (A) proportion of grass cover constituting grass taxa endemic to the DMC per plot; (B) proportion of grass cover constituting alien grass taxa per plot (see Table S2 for taxa); (C) mean range size, calculated as alpha hull=2, of grass taxa endemic to sub-Saharan Africa per plot. Boxplots are colour coded according to grazing regime, with colour codes of grazing and burning regimes found on the x axis as well as number of 2×2m plots (N) also indicated. Different letters above boxplots denote significant (p <0.05) differences following two-way ANOVA on GLM results and post-hoc Tukey test.

**Figure S3.**
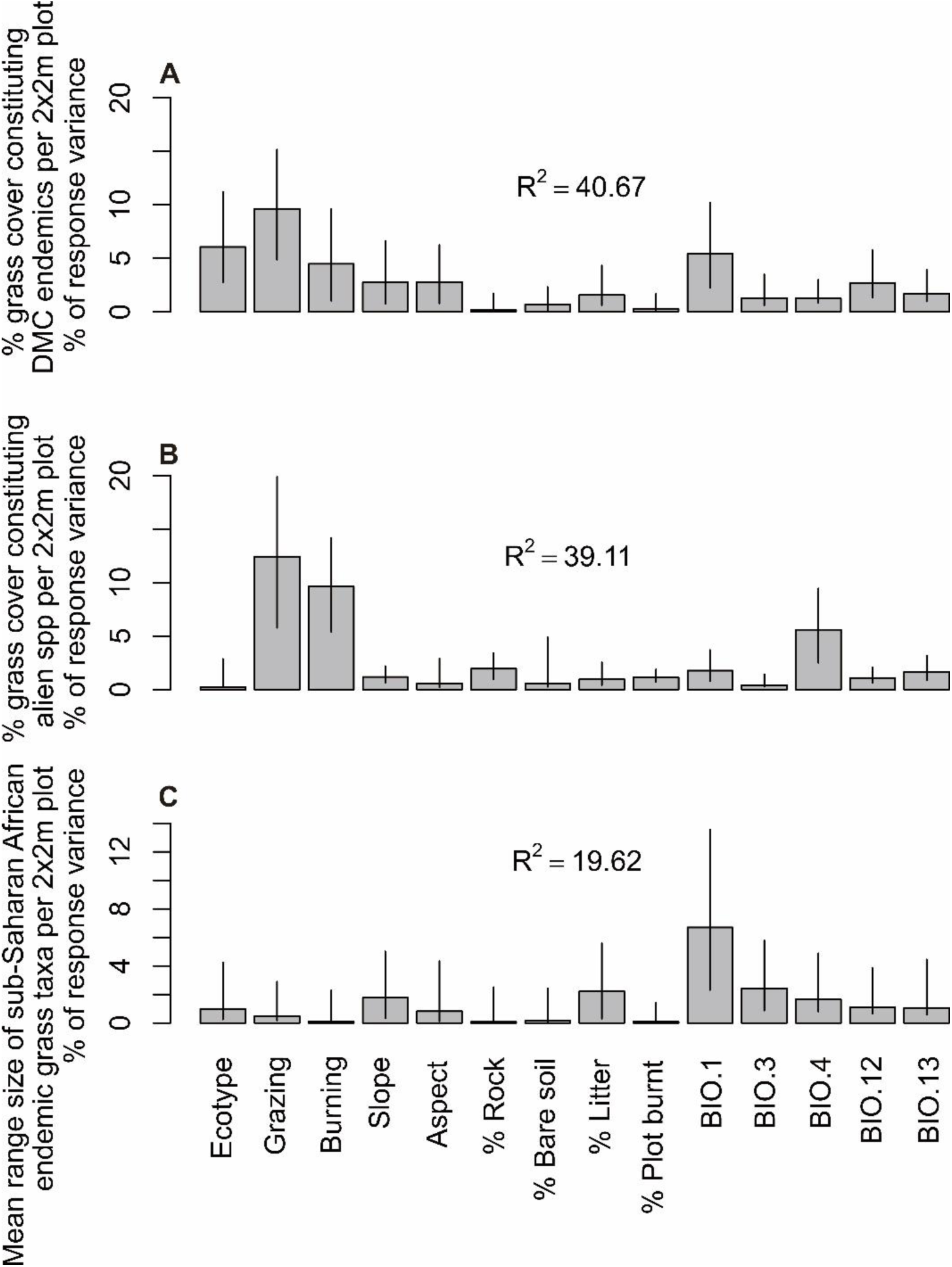
Relative importance of environmental vectors on plant composition and range size traits (A-C). Shows % response of variance and 95% confidence intervals of environmental vectors when analysing (A) proportion of grass cover constituting grass taxa endemic to the DMC per plot; (B) proportion of grass cover constituting alien grass taxa per plot (see Table S2 for taxa); (C) mean range size, calculated as alpha hull=2, of grass taxa endemic to sub-Saharan Africa per plot. Values based on relative importance analysis using the lmg method. R^2^ values of each analysis are shown within plots. Description of BioClim variable codes found in Table S1.

**Figure S4.**
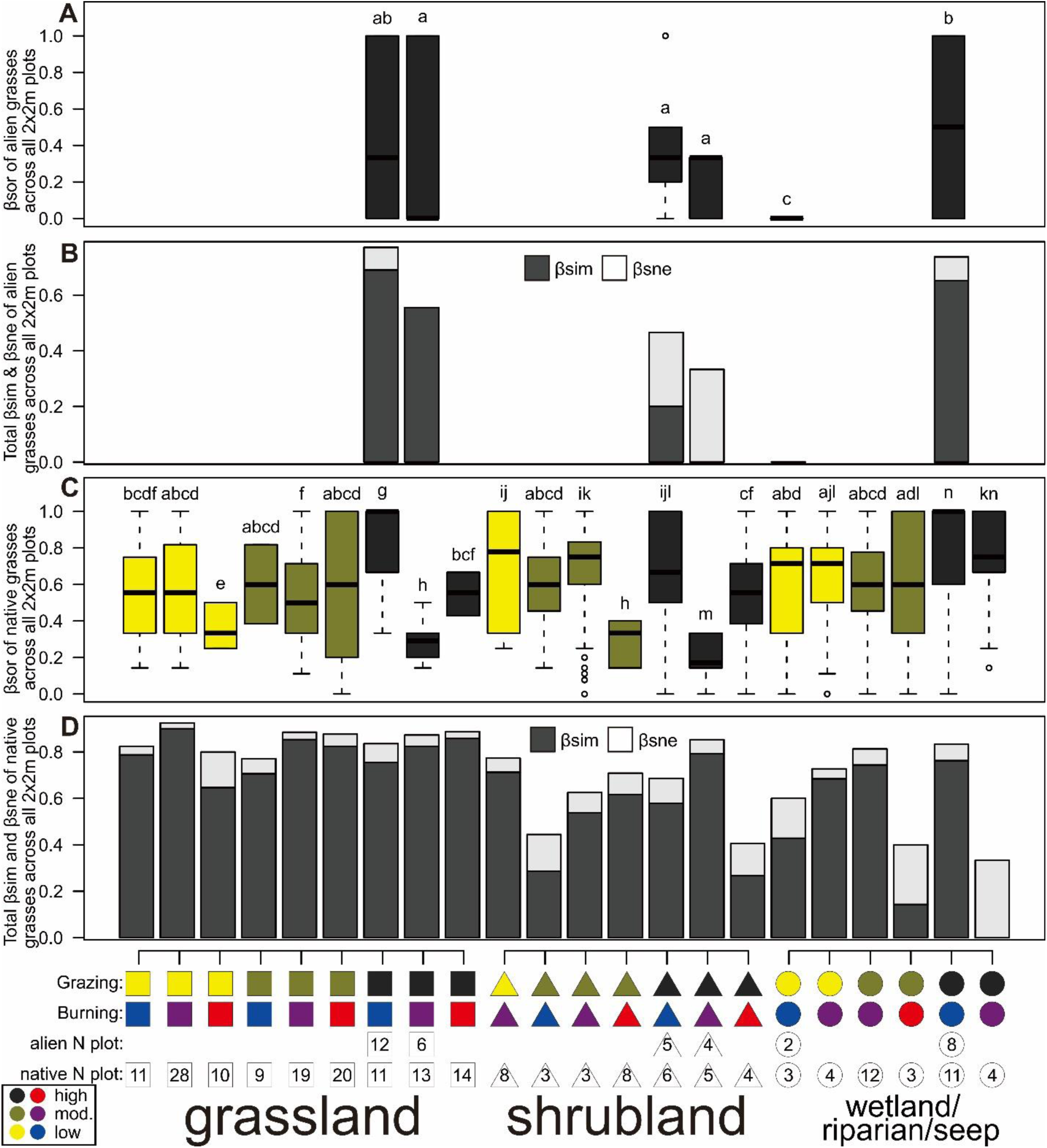
Alien (A-B) and native (C-D) grass community Beta diversity traits across different ecotype and disturbance regimes at a plot-level. Comparisons of (A) plot-level beta diversity of alien grass taxa, represented by overall Sørensen dissimilarity (βsor) of ecotype-disturbance categories across 2×2m plots; (B) contribution of turnover (βsim) and nestedness (βsne) to overall βsor for alien grasses; (C) plot-level beta diversity of native grass taxa, represented by overall βsor of ecotype-disturbance categories across 2×2m plots; (D) contribution of βsim and βsne to overall βsor for native grasses. Boxplots of (A) and (C) are colour coded according to grazing regime, with colour codes of grazing and burning regimes found on the x axis as well as sample size i.e. number of 2×2m plots for analysis of alien (alien N plot) or native (native N plot) grasses. Different letters above boxplots of (A) and (C) denote significant (p <0.05) differences following two-way ANOVA on binomial GLM results and post-hoc Tukey test.

**Table S1.** WorldClim bioclimatic (BioClim) variable codes and description

| Code | Description |
| --- | --- |
| BIO.1 | **annual mean temperature** |
| BIO.2 | **mean diurnal range (mean of monthly (max temp - min temp))** |
| BIO.3 | **isothermality (BIO.2/BIO.7) (×100)** |
| BIO.4 | **temperature seasonality (standard deviation ×100)** |
| BIO.5 | **max temperature of warmest month** |
| BIO.6 | **min temperature of coldest month** |
| BIO.7 | **temperature annual range (BIO.5-BIO.6)** |
| BIO.8 | **mean temperature of wettest quarter** |
| BIO.9 | **mean temperature of driest quarter** |
| BIO.10 | **mean temperature of warmest quarter** |
| BIO.11 | **mean temperature of coldest quarter** |
| BIO.12 | **annual precipitation** |
| BIO.13 | **precipitation of wettest month** |
| BIO.14 | **precipitation of driest month** |
| BIO.15 | **precipitation seasonality (coefficient of variation)** |
| BIO.16 | **precipitation of wettest quarter** |
| BIO.17 | **precipitation of driest quarter** |
| BIO.18 | **precipitation of warmest quarter** |
| BIO.19 | **precipitation of coldest quarter** |

**Table S2.** DMC endemic [according to Carbutt (2019), Linder et al. (2014), and Sylvester et al. (2020)] and alien grass taxa encountered in plots [according to Fish et al. (2015), Soreng et al. (2020), and Sylvester et al. (in press)]

| DMC endemic | Alien |
| --- | --- |
| *Catabrosa drakensbergensis* (Hedberg & I. Hedberg) Soreng & Fish | *Agrostis gigantea* Roth |
| *Festuca caprina* var. *irrasa* Stapf | *Bromus catharticus* Vahl |
| *Festuca caprina* var. *macra* Stapf | *Deschampsia cespitosa* (L.) P. Beauv. |
| *Festuca drakensbergensis* Sylvester, Soreng & M.D.P.V. Sylvester | *Festuca rubra* L. |
| *Pentameris exserta* (H.P. Linder) Galley & H.P. Linder | *Poa annua* L. |
| *Tenaxia disticha* var. *dracomontana* H.P. Linder | *Poa compressa* L. |
| *Tenaxia disticha* var. *lustricola* H.P. Linder | *Poa pratensis* subsp. *irrigata* (Lindm.) H. Lindb. |
| *Trisetopsis galpinii* (Schweick.) Röser & A. Wölk | *Poa pratensis* subsp. *pratensis* L. |

**Table S3.** Weightings on CCA axis 1 and 2, R^2^ and significance values of environmental vectors plotted onto CCA plot. Signif. codes: 0 ‘***’ 0.001 ‘**’ 0.01 ‘*’.

| Vector | CCA1 | CCA2 | r2 | Pr(>r) |
| --- | --- | --- | --- | --- |
| Grazing regime | 0.98347 | -0.18106 | 0.2976 | 0.001 *** |
| Burning regime | -0.94784 | 0.31875 | 0.2355 | 0.001 *** |
| Slope | -0.58659 | 0.80988 | 0.3382 | 0.001 *** |
| Aspect | -0.55671 | 0.8307 | 0.1581 | 0.001 *** |
| % Rock | -0.98858 | 0.1507 | 0.0448 | 0.009 ** |
| % Bare soil | 0.96683 | 0.25543 | 0.0686 | 0.002 ** |
| % Litter | -0.96252 | 0.2712 | 0.0731 | 0.002 ** |
| % Plot burnt | -0.93391 | -0.35751 | 0.0354 | 0.033 * |
| BIO.1 | 0.48879 | 0.8724 | 0.6253 | 0.001 *** |
| BIO.3 | 0.2634 | 0.96469 | 0.3004 | 0.001 *** |
| BIO.4 | 0.9996 | 0.02831 | 0.1753 | 0.001 *** |
| BIO.12 | -0.44978 | -0.89314 | 0.2296 | 0.001 *** |
| BIO.13 | -0.99836 | 0.0572 | 0.1491 | 0.001 *** |

**Table S4.**
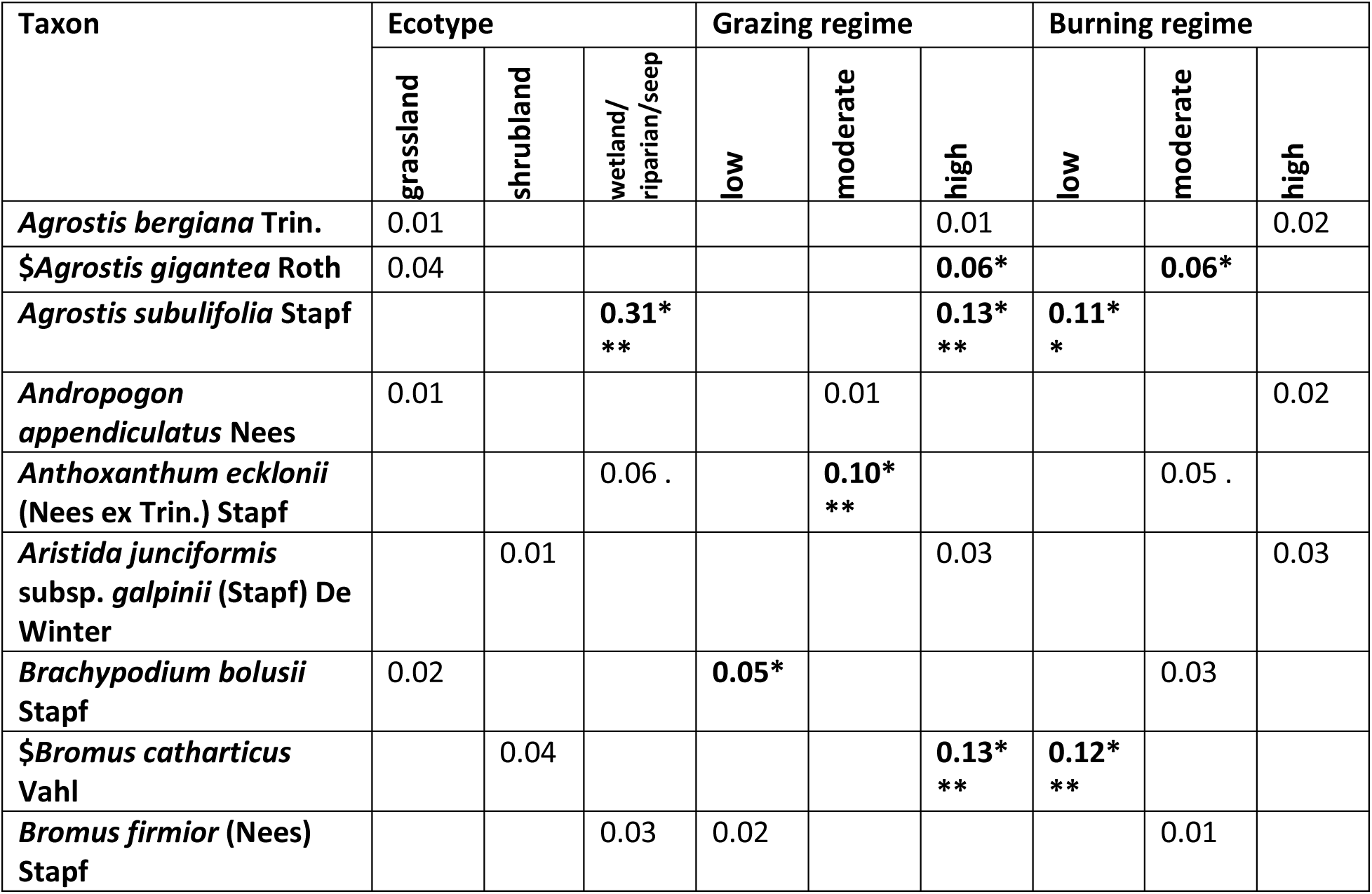
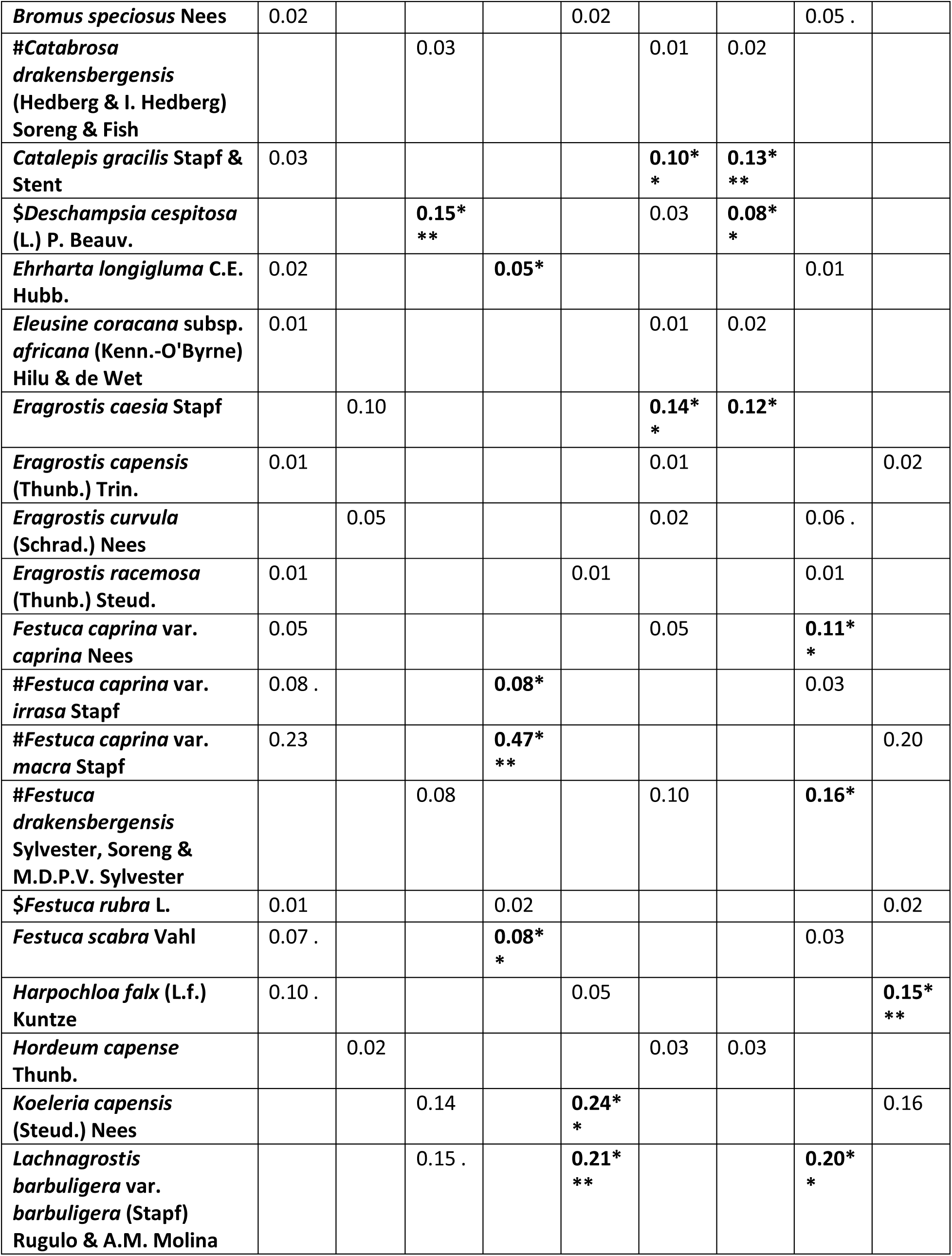
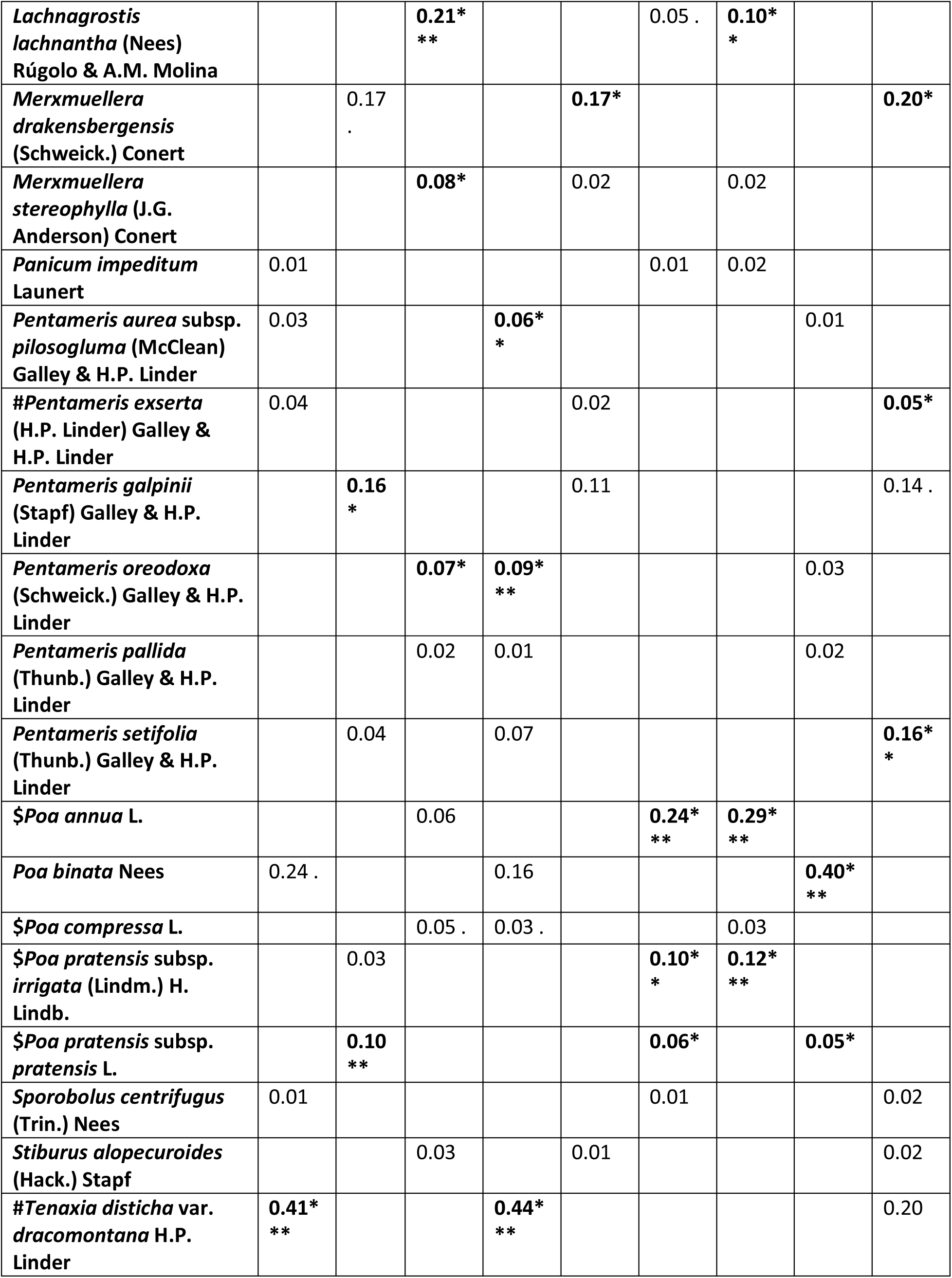
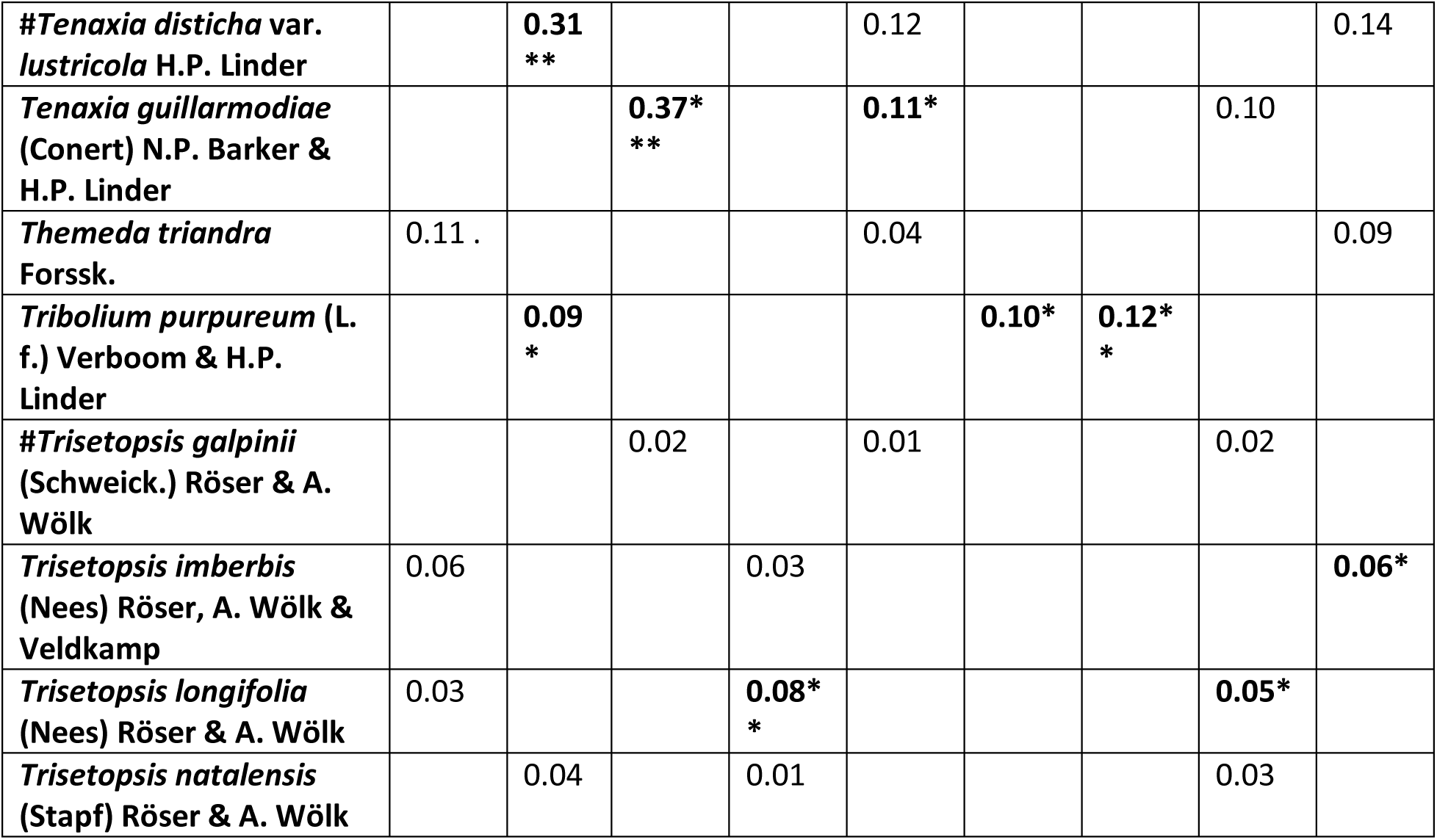
Indval values for different grass taxa after performing three separate indicator species analyses comparing: (1) ecotypes, (2) grazing regimes, and (3) burning regimes. Significant (p <0.05) Indval values are in bold. Signif. codes: ‘***’=p<0.001, ‘**’=p<0.01,’*’=p<0.05, ‘.’=p<0.1. Alien and DMC endemic taxa are denoted with a ‘$’ or ‘#’, respectively.

## Notes

### Competing Interest Statement

The authors have declared no competing interest.

## References

Adelabu, D. B., Clark, V. R., & Bredenhand, E. (2020). Potential for Sustainable Mountain Farming: Challenges and Prospects for Sustainable Smallholder Farming in the Maloti– Drakensberg Mountains. Mountain Research and Development 40(*1*), A1–A11. https://doi.org/10.1659/MRD-JOURNAL-D-19-00058.1

Adie, H., Kotze, D. J., & Lawes, M. J. (2017). Small fire refugia in the grassy matrix and the persistence of Afrotemperate forest in the Drakensberg mountains. Scientific Reports-UK, 7, 1–10.

Archibald, S., Staver, C., & Levin, S. A. (2012). Evolution of human-driven fire regimes in Africa. Proceedings of the National Academy of Sciences of the United States of America, 109, 847–852. https://doi.org/10.1073/pnas.1118648109.

Barker, N. P., & Fish, L. (2007). Rare and infrequent southern African grasses: assessing their conservation status and understanding their biology. Biodiversity and Conservation, 16, 4051–4079.

Baselga, A., Orme, D., Villeger, S., De Bortoli, J., Leprieur, F., & Logez, M. (2020). betapart: Partitioning Beta Diversity into Turnover and Nestedness Components. R package version 1.5.2. https://CRAN.R-project.org/package=betapart

Bentley, L. K., & O’Connor, T. G. (2018). Temperature control of the distributional range of five C3 grass species in the uKhahlamba-Drakensberg Park, KwaZulu-Natal, South Africa, African Journal of Range & Forage Science, 35(*1*), 45–54. https://doi.org/10.2989/10220119.2018.1459841.

Bentley, L. K., Robertson, M. P., & Barker, N. P. (2019). Range contraction to a higher elevation: The likely future of the montane vegetation in South Africa and Lesotho. Biodiversity and Conservation, 28(*1*), 131–153. https://doi.org/10.1007/s10531-018-1643-6

Binkley, D., & Fisher, R. F. (2019). Ecology and Management of Forest Soils, 5th Edition. Wiley-Blackwell.

Bond, W. J., Stevens, N., Midgley, G. F., & Lehmann, C. E. R. (2019). The Trouble with Trees: Afforestation Plans for Africa. Trends in Ecology & Evolution, 34(*11*), 963–965. https://doi.org/10.1016/j.tree.2019.08.003

Borer, E., Seabloom, E., Gruner, D. et al. (2014). Herbivores and nutrients control grassland plant diversity via light limitation. Nature, 508, 517–520. https://doi.org/10.1038/nature13144

Brand, R. F., Scott-Shaw, C. R., & O’Connor, T. G. (2019). The alpine flora on inselberg summits in the Maloti–Drakensberg Park, KwaZuluNatal, South Africa. Bothalia, 49(*1*), a2386. https://doi.org/10.4102/abc.v49i1.23

Breman, E., Ekblom, A., Gillson, L., & Norström, E. (2019). Phytolith-based environmental reconstruction from an altitudinal gradient in Mpumalanga, South Africa, 10,600 BP– present. Review of Palaeobotany and Palynology, 263, 104–116. https://doi.org/10.1016/j.revpalbo.2019.01.001.

Carbutt, C. (2019). The Drakensberg Mountain Centre: A necessary revision of southern Africa’s high-elevation centre of plant endemism. South African Journal of Botany, 124, 508– 529.

Carbutt, C. (2020). The Imperiled Alpine Grasslands of the Afrotropic Realm. Reference Module in Earth Systems and Environmental Sciences. Elsevier. https://doi.org/10.1016/B978-0-12-821139-7.00012-X

Carbutt, C., & Edwards, T. J. (2004). The flora of the Drakensberg Alpine Centre. Edinburgh Journal of Botany, 60, 581–607.

Carbutt, C., & Edwards, T. J. (2006). The endemic and near-endemic angiosperms of the Drakensberg Alpine Centre. South African Journal of Botany, 72, 105–132.

Carbutt, C., & Edwards, T. J. (2015). Reconciling ecological and phytogeographical spatial boundaries to clarify the limits of the montane and alpine regions of sub-Sahelian Africa. South African Journal of Botany, 98, 64–75. https://doi.org/10.1016/j.sajb.2015.01.014.

Dauby, G. (2018). ConR: Computation of parameters used in preliminary assessment of conservation status. R package version 1.2.2. https://CRAN.R-roject.org/package5ConR.

Dufrêne, M., & Legendre, P. (1997). Species assemblages and indicator species: the need for a flexible asymmetrical approach. Ecological Monographs, 67, 345–366.

Ficetola, G. F., Rondinini, C., Bonardi, A., Katariya, V., Padoa-Schioppa, E., & Angulo, A. (2014). An evaluation of the robustness of global amphibian range maps. Journal of Biogeography, 41, 211–221. https://doi.org/10.1111/jbi.12206.

Finch, J. M., Hill, T. R., Meadows, M. E., Lodder, J., & Bodmann, L. (In press). Fire and montane vegetation dynamics through successive phases of human occupation in the northern Drakensberg, South Africa. Quaternary International. https://doi.org/10.1016/j.quaint.2021.01.026.

Fish, L., Mashau, A. C., Moeaha, M. J., & Nembudani, M. T. (2015). Identification guide to southern African grasses: an identification manual with keys, descriptions and distributions. Strelizia, 36. South African National Biodiversity Institute.

Gehrke, B., & Linder, H. P. (2014). Species richness, endemism and species composition in the tropical Afroalpine flora. Alpine Botany, 124, 165–177.

Goslee, S. C., & Urban, D. L. (2007). The ecodist package for dissimilarity-based analysis of ecological data. Journal of Statistical Software, 22(*7*), 1–19.

Grab, S. W., & Nash, D. J. (In press) “But what silence! No more gazelles…”: Occurrence and extinction of fauna in Lesotho, southern Africa, since the late Pleistocene. Quaternary International. https://doi.org/10.1016/j.quaint.2020.04.030.

Grab, S., Knight, J., Mol, L., Botha, T., Carbutt, C., & Woodborne, S. (2021). Periglacial landforms in the high Drakensberg, Southern Africa: morphogenetic associations with rock weathering rinds and shrub growth patterns, Geografiska Annaler: Series A, Physical Geography. https://doi.org/10.1080/04353676.2020.1856625

Grömping, U. (2006) Relative Importance for Linear Regression in R: The Package relaimpo. Journal of Statistical Software, 17(*1*), 1–27. https://doi.org/10.18637/jss.v017.i01

Harrell, F. E. Jr, with contributions from Charles Dupont and many others (2020). Hmisc: Harrell Miscellaneous. R package version 4.4-1. https://CRAN.R-project.org/package=Hmisc

Johansson, M., Fetene, M., Malmer, A., & Granström, A. (2012). Tending for cattle: traditional fire management in Ethiopian Montane Heathlands. Ecology and Society, 17(*3*), 19. http://dx.doi.org/10.5751/ES-04881-170319

Johansson, M. U., Frisk, C. A., Nemomissa, S., & Hylander, K. (2018). Disturbance from traditional fire management in subalpine heathlands increases Afro-alpine plant resilience to climate change. Global Change Biology, 24, 2952–2964. https://doi.org/10.1111/gcb.14121

Joubert, L., Pryke, J. S., & Samways, M. J. (2017). Moderate grazing sustains plant diversity in Afromontane grassland. Applied Vegetation Science, 20, 340–351. https://doi.org/10.1111/avsc.12310

Killick, D. J. B. (1994). Drakensberg Alpine Region—Lesotho and South Africa. In S. D. Davis, V. H. Heywood, A. C. Hamilton (Eds.), Centres of Plant Diversity—A Guide and Strategy for Their Conservation (pp. 257–260). IUCN Publications Unit, Cambridge.

Knight, J., & Grab, S. W. (2015). The Drakensberg escarpment: Mountain processes at the edge. In S. W. Grab & J. Knight (Eds.), Landscapes and Landforms of South Africa (pp. 47–55). Springer.

Körner, C. (2003). Alpine Plant Life - Functional Plant Ecology of High Mountain Ecosystems, 2nd edition. Springer.

Körner, C. (2012). Alpine Treelines: Functional Ecology of the Global High Elevation Tree Limits. Springer.

Lehmann, C. E., Archibald, S. A., Hoffmann, W. A., & Bond, W. J. (2011). Deciphering the distribution of the savanna biome. New Phytologist, 191, 197–209. https://doi.org/10.1111/j.1469-8137.2011.03689.x

Lézine, A. -M., Izumi, K., Kageyama, M., & Achoundong, G. (2019). A 90,000-year record of Afromontane forest responses to climate change. Science, 363, 177–181.

Lindeman, R. H., Merenda, P. F., & Gold, R. Z. (1980). Introduction to Bivariate and Multivariate Analysis. Scott, Foresman, Glenview, IL.

Linder, H. P., Perl, F., Bouchenak-Khelladi, Y., & Barker, N. P. (2014). Phylogeographical Pattern in the Southern African Grass *Tenaxia disticha* (Poaceae). Systematic Botany, 39, 428– 440. https://doi.org/10.1600/036364414X680906

Lodder, J., Hill, T. R., & Finch, J. M. (2018). A 5000-yr record of Afromontane vegetation dynamics from the Drakensberg Escarpment, South Africa. Quaternary International, 470, 119–129. https://doi.org/10.1016/j.quaint.2017.08.019

Mahibbur, M. R., & Govindarajulu, Z. (1997). A modification of the test of Shapiro and Wilk for normality. Journal of Applied Statistics, 24, 219–236. https://doi.org/10.1080/02664769723828

McKinney, M. L., & Lockwood, J. L. (1999). Biotic homogenizaton: a few winners replacing many losers in the next mass extinction. Trends in Ecology and Evolution, 14, 450–453.

Meadows, M., & Linder, P. J. (1993). A Palaeoecological Perspective on the Origin of Afromontane Grasslands. Journal of Biogeography, 20, 345–355.

Morris, C., Everson, C., Everson, T. M., & Gordijn, P. (2021). Frequent burning maintained a stable grassland over four decades in the Drakensberg, South Africa. African Journal of Range & Forage Science, 38, 39–52. https://doi.org/10.2989/10220119.2020.1825120.

Mucina, L., & Rutherford, M. C. (2006). The Vegetation of South Africa, Lesotho and Swaziland. South African National Biodiversity Institute.

Ngwenya, S. J., Torquebiau, E., & Ferguson, J. W. H. (2019). Mountains as a critical source of ecosystem services: the case of the Drakensberg, South Africa. Environment, Development and Sustainability, 21, 1035–1052. https://doi.org/10.1007/s10668-017-0071-1

O’Connor, T. G., Kuyler, P., Kirkman, K. P., & Corcoran, B. (2010). Which grazing management practices are most appropriate for maintaining biodiversity in South African grassland? African Journal of Range & Forage Science, 27(*2*), 67–76. https://doi.org/10.2989/10220119.2010.502646

Oksanen, J., Blanchet, F. G., Friendly, M., Kindt, R., Legendre, P., McGlinn, D., Minchin, P. R., O’Hara, R. B., Simpson, G. L., Solymos, P., Stevens, M. H. H., Szoecs, E., & Wagner, H. (2019). vegan: Community Ecology Package. R package version 2.5-6. https://CRAN.R-project.org/package=vegan.

Olden, J. D., Comte, L., & Giam, X. (2018). The Homogocene: a research prospectus for the study of biotic homogenisation. NeoBiota, 37, 23–36. https://doi.org/10.3897/neobiota.37.22552

Padgham, M., & Sumner, M. D. (2021). geodist: Fast, Dependency-Free Geodesic Distance Calculations. R package version 0.0.7. https://CRAN.R-project.org/package=geodist

Plants Of The World Online (2021) http://www.plantsoftheworldonline.org [accessed 15.2.2021]

R Core Team (2020) R: A language and environment for statistical computing. R Foundation for Statistical Computing, Vienna, Austria. https://www.R-project.org/.

Ripley, B., Venables, B., Bates, D. M., Hornik, K., Gebhardt, A., & Firth, D. (2020). MASS Version:7.3-53: Support Functions and Datasets for Venables and Ripley’s MASS. https://cran.r-project.org/web/packages/MASS/index.html.

Roberts, D. W. (2019). labdsv: Ordination and Multivariate Analysis for Ecology. R package version 2.0-1. https://CRAN.R-project.org/package=labdsv

Smouse, P., Long, J., & Sokal, R. (1986). Multiple regression and correlation extensions of the Mantel test of matrix correspondence. Systematic Zoology, 35, 627–632.

Soreng, R. J., Sylvester, S. P., Sylvester, M. D. P. V., & Clark, V. R. (2020). New records and key to *Poa* (Pooideae: Poaceae) from the Flora of Southern Africa region, and notes on taxa including a diclinous breeding system in *Poa binata*. PhytoKeys, 165, 27–50. https://doi.org/10.3897/phytokeys.165.55948

Stein, A., Gerstner K., & Kreft H. (2014). Environmental heterogeneity as a universal driver of species richness across taxa, biomes and spatial scales. Ecology Letters, 17, 866–880. https://doi.org/10.1111/ele.12277

Stewart, B. A., Parker, A. G., Dewar, G., Morley, M. W., & Allott, L. F. (2016). Chapter 14: Follow the Senqu: Maloti-Drakensberg Paleoenvironments and Implications for Early Human Dispersals into Mountain Systems. In S. C. Jones & B. A. Stewart (Eds.), Africa from MIS 6-2: Population Dynamics and Paleoenvironments, Vertebrate Paleobiology and Paleoanthropology (pp. 247–271). Springer. https://doi.org/10.1007/978-94-017-7520-5_14

Strömberg, C. A. (2011). Evolution of grasses and grassland ecosystems. Annual Review of Earth and Planetary Sciences, 39, 517–544.

Sylvester, S. P., Sylvester, M. D. P. V., & Kessler, M. (2014). Inaccessible ledges as refuges for the natural vegetation of the high Andes. Journal of Vegetation Science, 25(*5*), 1225–1234. http://dx.doi.org/10.1111/jvs.12176

Sylvester, S. P., Heitkamp, F., Sylvester, M. D. P. V., Jungkunst, H. F., Sipman, H. J. M., Toivonen, J. M., Gonzales Inca, C., Ospina González, J. C., & Kessler, M. (2017). Relict high-Andean ecosystems challenge our concepts of naturalness and human impact. Scientific Reports-UK, 7, 3334. http://dx.doi.org/10.1038/s41598-017-03500-7

Sylvester, S. P., Soreng, R. J., Sylvester, M. D. P. V., & Clark, V. R. (2020). *Festuca drakensbergensis* (Poaceae): A common new species in the *F. caprina* complex from the Drakensberg Mountain Centre of Floristic Endemism, southern Africa, with key and notes on taxa in the complex including the overlooked *F. exaristata*. PhytoKeys, 162, 45– 69. https://doi.org/10.3897/phytokeys.162.55550

Sylvester, S. P., Soreng, R. J., Sylvester, M. D. P. V., Mapaura, A., & Clark, V. R. (In press). New records of exotic and potentially invasive grass (Poaceae) species for southern Africa. Bothalia.

Tamme, R., Hiiesalu, I., Laanisto, L., Szava-Kovats, R., & Pärtel, M. (2010). Environmental heterogeneity, species diversity and co-existence at different spatial scales. Journal of Vegetation Science, 21, 796–801. https://doi.org/10.1111/j.1654-1103.2010.01185.x

Taylor, S. J., Ferguson, J. W. H., Engelbrecht, F. A., Clark, V. R., Van Rensburg, S., & Barker, N. (2016). The Drakensberg escarpment as the great supplier of water to South Africa. In G. B. Greenwood & J. F. Schroder (Eds.), Mountain Ice and Water (pp. 1–46). Elsevier. https://doi.org/10.1016/B978-0-444-63787-1.00001-9.

Ter Braak, C. J. F. (1986). Canonical correspondence analysis: a new eigenvector technique for multivariate direct gradient analysis. Ecology, 67(*5*), 1167–1179.

Thiers, B. (2021). Index Herbariorum: A global directory of public herbaria and associated staff. New York Botanical Garden’s Virtual Herbarium. http://sweetgum.nybg.org/ih/ [accessed 5 May 2020].

Tonidandel, S., & LeBreton, J. M. (2011). Relative Importance Analysis: A Useful Supplement to Regression Analysis. The Journal of Business and Psychology, 26, 1–9. https://doi.org/10.1007/s10869-010-9204-3

Turpie, J., Benn, G., Thompson, M., & Barker, N. (2021). Accounting for land cover changes and degradation in the Katse and Mohale Dam catchments of the Lesotho highlands. African Journal of Range and Forage Science, 38, 53–66. https://doi.org/10.2989/10220119.2020.1846214

Van Coller, H., Siebert, F., Scogings, P. F., & Ellis, S. (2018). Herbaceous responses to herbivory, fire and rainfall variability differ between grasses and forbs. South African Journal of Botany, 119, 94–103. https://doi.org/10.1016/j.sajb.2018.08.024

Van Wyk, A. E., & Smith, G. F. (2001). Regions of floristic endemism in southern Africa. Umdaus.

Vidal, J. D., & Clark, V. R. (2020). Afro-Alpine Plant Diversity in the Tropical Mountains of Africa. In M. I. Goldstein & D. A. Della Sala (Eds.) Encyclopedia of the World’s Biomes (pp. 373– 394). Elsevier. https://doi.org/10.1016/B978-0-12-409548-9.11885-8

Wei, T., & Simko, V. (2017). R package “corrplot”: Visualization of a Correlation Matrix (Version 0.84). https://github.com/taiyun/corrplot

Wesche, K., Miehe, G., & Kaeppeli, M. (2000). The significance of fire for Afroalpine ericaceous vegetation. Mountain Research and Development, 20, 340–347.

White, F. (1983). The vegetation of Africa. UNESCO, Paris.

Wood, J. R. I. (1997). A Handbook of the Yemen Flora. The Board of Trustees of the Royal Botanic Gardens, Kew

Zaloumis, N. P., & Bond, W. J. (2016). Reforestation or conservation? The attributes of old growth grasslands in South Africa. Philosophical Transactions of the Royal Society, B, 371(*1703*), 20150310. http://dx.doi.org/10.1098/rstb.2015.0310

Zizka, A., Silvestro, D., Andermann, T., Azevedo, J., Duarte Ritter, C., Edler, D., Farooq, H., Herdean, A., Ariza, M., Scharn, R., Svantesson, S., Wengström, N., Zizka, V., & Antonelli, A. (2019). CoordinateCleaner: Standardized cleaning of occurrence records from biological collection databases. Methods in Ecology and Evolution, 10, 744–751. http://dx.doi.org/10.1111/2041-210X.13152.

